# Genome-wide exploration of the transcriptional regulatory landscape in the early-diverging fungus *R. microsporus* reveals pervasive DNA methyl adenine regulatory network

**DOI:** 10.1101/2025.01.24.634581

**Authors:** Carlos Lax, Leo A. Baumgart, Ghizlane Tahiri, Natalia Nicolás-Muñoz, Yu Zhang, Ian K. Blaby, Ronan C O’Malley, Vivian Ng, Eusebio Navarro, Igor V Grigoriev, Francisco E. Nicolás, Victoriano Garre

**Affiliations:** Departamento de Genética y Microbiologá, Facultad de Biologá, Universidad de Murcia, 30100 Murcia, Spain; U.S. Department of Energy Joint Genome Institute, Lawrence Berkeley National Laboratory, Berkeley, CA 94720 USA; Department of Human Genetics, University of Chicago, Chicago, IL, USA; Environmental Genomics and Systems Biology Division, Lawrence Berkeley National Laboratory, Berkeley, CA, 94720, USA; Department of Plant and Microbial Biology, University of California Berkeley, Berkeley, CA 94720 USA

## Abstract

Genetic regulation mechanisms rely on complex transcriptional networks that are often difficult to decipher. The study of transcription factor (TF) binding sites and their targets has traditionally faced scalability challenges, hindering comprehensive cistrome analyses. However, the development of the DNA affinity purification and sequencing (DAP-seq) technique has allowed unprecedented large-scale studies at genome-wide level of TF binding with high reproducibility. In this study, we apply this technique to the human opportunistic pathogen *R. microsporus*, a mucoralean fungus belonging to the understudied group of early-diverging fungi (EDF). We characterize genome-wide binding sites of 58 TFs encoded by genes regulated through adenine methylation and representing major TF families, representing the most extensive DAP-seq study in filamentous fungi. This analysis reveals their binding profiles and recognized sequences, expanding and diversifying the catalog of known fungal motifs. By integrating this data with DNA 6-methyladenine profiling, we uncover the extensive direct and indirect impact of this epigenetic modification on the regulation of gene expression. Furthermore, the generated data facilitates the identification and functional characterization of TFs involved in biologically relevant processes, such as zinc metabolism and light response, serving as a proof of concept for the utility of the DAP-seq data. These findings not only enhance our understanding of regulatory mechanisms in *R. microsporus* but also provide broader insights into gene regulation across the fungal kingdom.

## Introduction

Transcription factors (TFs) play a central role in gene expression regulation by binding to specific DNA sequences. They can either activate or repress the transcription of target genes, thereby modulating cellular processes in response to various environmental and developmental signals. The complete set of genomic binding sites for a given TF, often referred to as its cistrome^1^, provides a comprehensive view of the complex transcriptional regulatory networks, which are also influenced by epigenetics modifications to ensure precise control of the transcription^2–4^.

Chromatin immunoprecipitation followed by sequencing (ChIP-seq) has been widely employed to determine TF binding sites (TFBS)^5–7^. However, scaling up this approach for genome-wide cistrome characterization is challenging due to its heavy reliance on antibody quality and availability, as well as its technical complexity when applying to proteins with low expression levels^8^. These limitations explain why TFBS are only available for specific proteins and are often limited to a few model organisms^6^. To overcome these issues, in vitro approaches that allow large-scale analyses of TSBS, such as protein binding microarrays (PBM)^9^ and systematic evolution of ligands by exponential enrichment (SELEX)^10^, were developed^11,12^. These methodologies, however, use synthetic short DNA fragments to identify the TF binding motif, which is then used to scan the genome for potential TFBS. The development of DNA affinity purification sequencing (DAP-seq), which combines the use of native genomic DNA of ChIP-seq and the scalability of in vitro approaches^13,14^, has enabled broad and comprehensive characterization of a considerable number of TFs from both prokaryotic and eukaryotic representatives^13,15,16^. Remarkably, this high-throughput methodology relies on the in vitro synthesis of the TFs of interest, followed by pulldown and next-generation sequencing of native or amplified genomic DNA, enabling cistrome characterization in a wide range of non-model organisms^15^.

Despite the development of these in vitro approaches, a significant gap remains in the study of TFBS in fungi. Fungi are an incredibly diverse kingdom of organisms with substantial medical, ecological, and industrial relevance^17–19^. Yet, compared to other eukaryotes, genome-wide studies of TF binding in fungi are very limited^20^. Most research has focused on characterizing specific TFs or DNA-binding proteins, predominantly in Dikarya fungi^21–23^, with only a few studies conducting broader analyses of TFBS across different TF families^24,25^. Noteworthy, since DAP-seq has made TFBS studies possible independent of species and the availability of molecular tools, it has been widely applied to plants^13,26–28^ and bacteria^15,16,29^, but only to some specific TFs on a smaller scale studies in fungi. In particular, this technique has allowed the elucidation of the fungal gene regulatory networks involved in nitrogen metabolism^30^ and carbon utilization^31^.

Beyond the Dikarya, the basal fungal subkingdom has been traditionally neglected. Combined with the limited availability of molecular tools, this has left many aspects of their biology, including gene expression regulation, largely unexplored. Furthermore, comprehensive studies on transcriptional regulators that respond to environmental signals, which are well-characterized in Dikarya fungi^30–33^, are notably lacking in early-diverging fungi (EDF). This group, however, encompasses the majority of known fungal phyla and presents a series of unique characteristics that make them of significant scientific interest^34,35^. Within this group, *Rhizopus microsporus* (Mucoromycota) stands out due to its high agricultural, industrial, and clinical relevance because of its phytopathogenic potential through symbiotic interaction with endobacteria^36,37^, the use of specific strains in the production of fermented products^38^, and its high frequency as a causal agent of mucormycosis, a lethal infection classified by the WHO as high-priority concern^39–41^. Notably, this fungus relies on the use of 6- methyladenine (6mA) as its main DNA epigenetic modification (Lax et al, under revision in Nature Communications), a unique characteristic shared with other EDF, ciliates, and green algae^42–44^. This modification is associated with gene expression in all these eukaryotic groups and its study in *R. microsporus* has revealed that it is essential (Lax et al, under revision in Nature Communications), underscoring its crucial role in gene regulation.

While most large-scale projects on cistrome analyses have focused on higher eukaryotes, such as mice and humans, fungal TF landscapes have received far less attention^20,45,46^. Here, we address this gap by presenting a comprehensive analysis of the *R. microsporus* cistrome. Using DAP-seq, we identified the binding sites of TFs regulated by 6mA through the use of DAP-seq, providing the most detailed gene regulatory network described in filamentous fungi to date. Moreover, a thorough investigation of TF families highly conserved across fungi, both phylogenetically close and distant relatives, offers valuable insights into the evolution of TFs and the mechanisms underlying gene regulation within the fungal kingdom.

## Results

### The fungal TF landscape

We curated a list of 51 TF families, including the fungal-specific families (APSES, MAT, and copper fist) based on previous studies^45,46^ and scanned the genomes of 62 fungal species, 43 of which are classified as EDF (**Supplementary Dataset 1**). In addition to the Zn cluster family expansion in Dikarya, particularly in Ascomycota^20^ (**Figure 1, Supplementary** Figure 1), other remarkable differences were found when comparing EDF representatives with Dikarya and among themselves. Contrary to the Zn cluster family, the HTH/Homeodomain-like family was more expanded in EDF compared to Ascomycota (p < 0.0001, Chi-square test) (**Figure 1, Supplementary** Figure 1), as is the C2H2/CCHC/CH/C5HC2 family in Monoblepharomycota, Neocallimastigomycota and Chytridiomycota (p < 0.0001, Chi-square test). Some TF families, such as E2F and Tubby, are absent in Dikarya^46^ (Figure 1, Supplementary Figure 1), while others, such as RFX and STAT, are good examples of TF families that are present in Dikarya but are absent or drastically less abundant in EDF.

**Figure 1.**
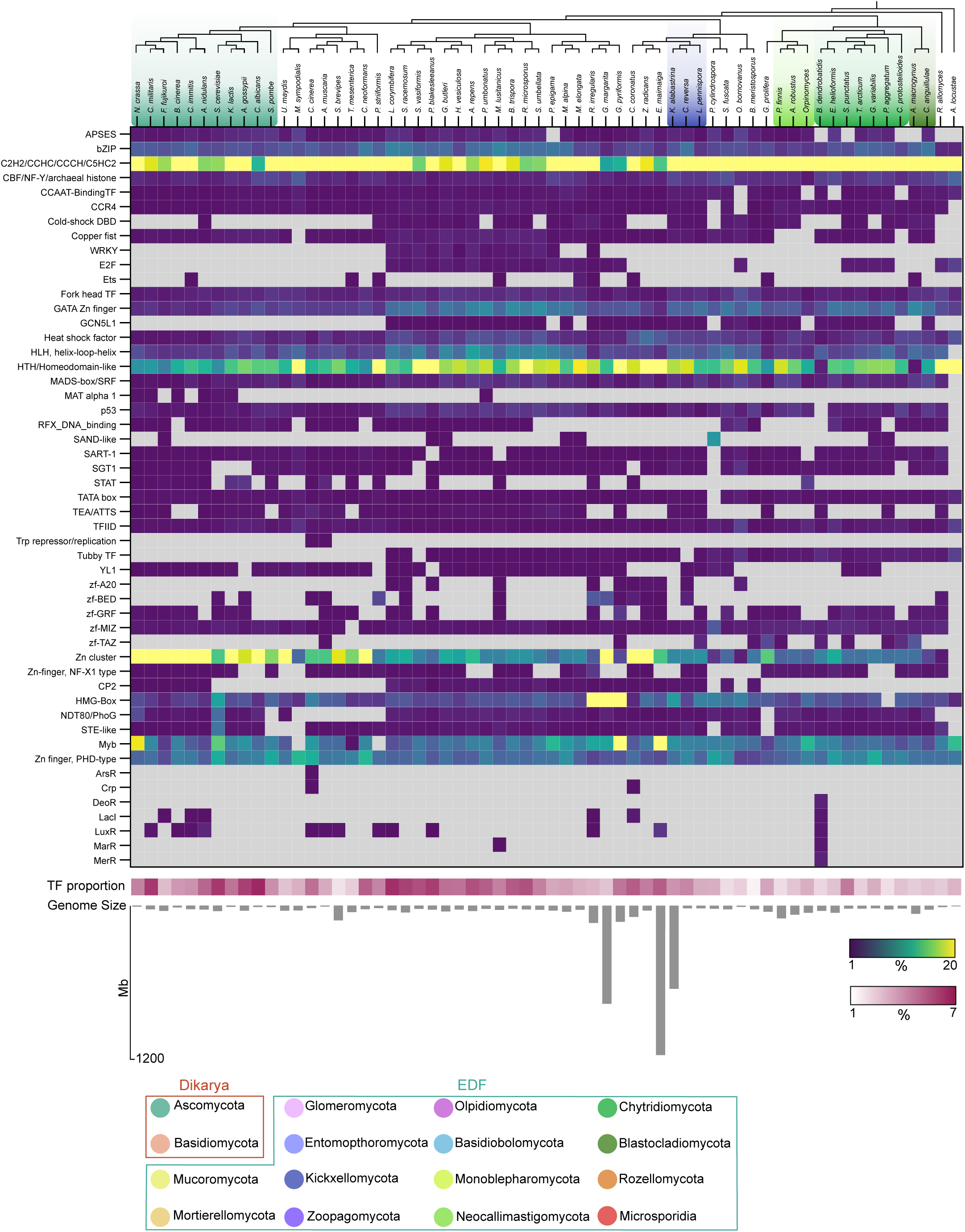
The fungal TF landscape. The abundance of each TF family, relative to the total number of TFs, is indicated for each species. Grey boxes indicate the absence of a given TF family. The bottom part displays the proportion of TF-coding genes relative to the total number of annotated genes, as well as the genome size, for each species. Information on the genomes analyzed can be found in Supplementary Dataset 1.

We also detected other interesting differences, such as the expansion of the HMG-box family in Glomeromycota, as well as the Helix-loop-helix (HLH) and GATA Zn finger families in Mucoromycota, Mortierellomycota, and some representatives of Chytridiomycota. Additionally, the Zn cluster TFs were particularly abundant in Entomopthoromycota (**Figure 1, Supplementary** Figure 1). This dynamic nature of TF families explains the notable expansion of TF coding genes, which is clearly noticeable in Ascomycota. Nevertheless, TF-coding genes also constitute a notable proportion of total protein-coding genes in Mucoromycota, Glomeromycota, and Entomopthoromycota (**Figure 1**). Noteworthy, Mucoromycota has the higher TF landscape diversity based on the Shannon index^46^, with some TF families, such as WRKY and E2F, being almost exclusively found in this phylum (**Figure 1**).

### Cistrome characterization of *R. microsporus* using DAP-seq

The diversity of TF landscapes found in Mucoromycota prompted us to investigate the cistrome of *R. microsporus*. For that purpose, we performed DAP-seq on a curated list of 88 TFs from major TF families (**Supplementary Table 1**) encoded by genes harboring 6mA concentrated in regions of high methylation density known as Methylated Adenine Clusters (MACs) under the same growth conditions used to generate DAP-seq samples (Lax et al. 2025, under revision in Nature Communications). MACs usually appear downstream of the transcription start site (TSS) of transcriptionally active genes of EDF (Mondo et al., 2017; Lax et al., 2024). We processed the datasets generated for each TF and retained only those with > 5% of the reads aligning to the predicted peaks for the respective TF (FRIP, Fragments of Reads in Peaks)^13,15^ (**Figure 2A and Supplementary Table 1**). Of the initial 88 TFs, 41 passed this filter, yielding a success ratio of 46.6%. To assess the data robustness and reproducibility, we conducted a second replicate of the experiment using the same TF candidates. In this replicate, 57 TFs passed the filter (64.7% success ratio), including 40 of the 41 TFs from the first attempt. In total, 58 TFs produced reliable results in at least one replicate and were considered for cistrome analysis. These TFs belonged to 11 different families, including the broadly distributed bZip, GATA Zn finger, HLH, Homeobox, and Heat-shock families (**Figure 2B**). Among these, TFs from the HLH, Homeobox, GATA, and HSF performed particularly well, with success rates of 89%, 92%, 82% and 87%, respectively (**Figure 2B**), whereas all TFs from CBF, Tubby, and PHD-type Zn finger failed to produce reliable data (**Figure 2B**). These results highlighted the influence of family-specific properties on success rates of DAP-seq experiments, as previously described^13^.

**Figure 2.**
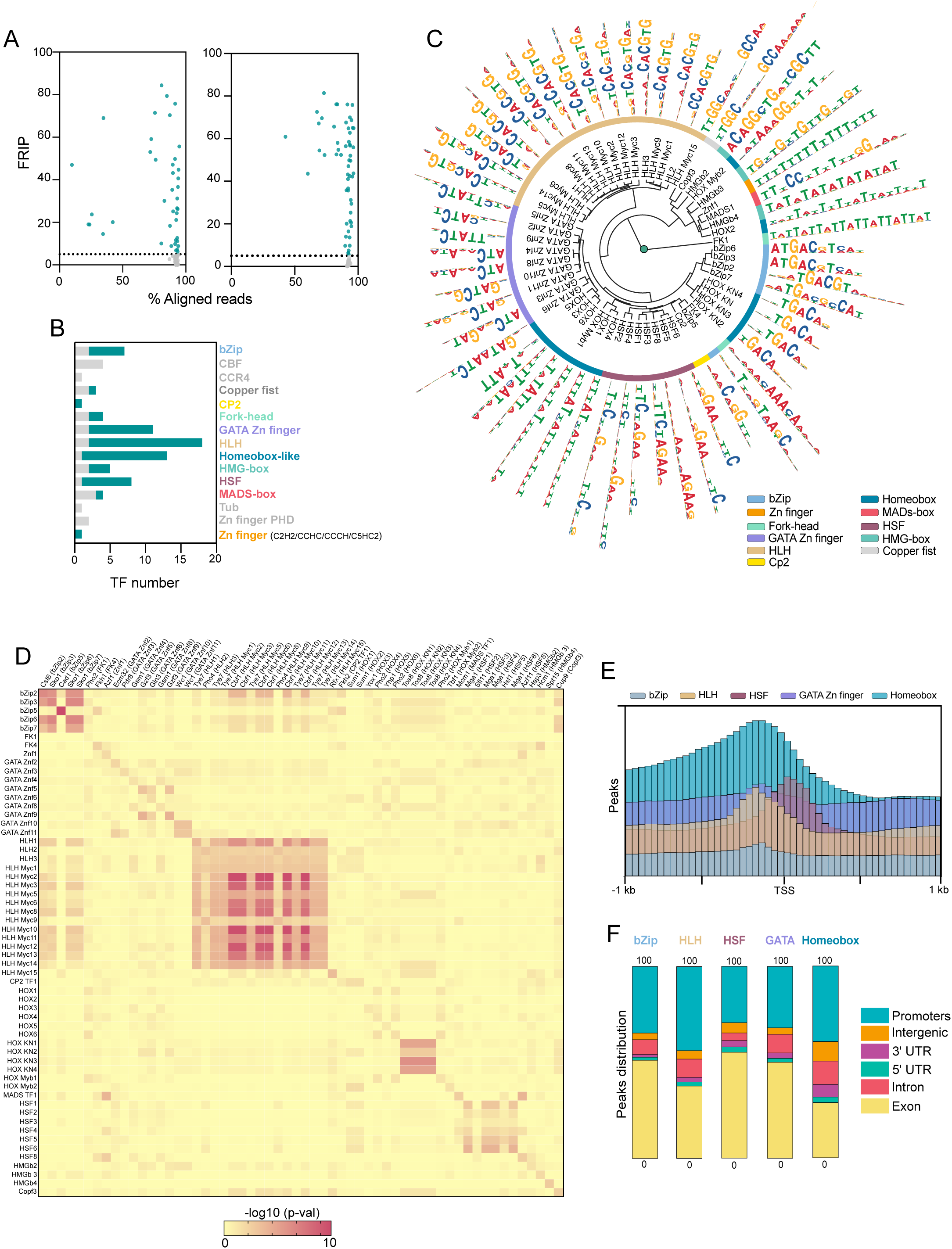
*R. microsporus* cistrome. (A) Filtering results for DAP-seq experiments indicating FRIP and the percentage of aligned reads. Each dot represents one TF. Only TFs with FRIP > 5% were considered. The left and the right panel are for the first and second replicate, respectively. (B). Proportion of TFs that passed filtering for each family. Grey color indicates FRIP < 5%. (C) TF motif clustering based on motif similarity using the R package motifStack. The outer circle indicates the TF family. (D) Comparison of *R. microsporus* DAP-seq motifs against JASPAR motif database. The best hit for each *R. microsporus* TF motif was compared against all new motifs identified. The color code represents the score (-log10 p-value) for each motif comparison. (E) Peak frequency distribution over protein-coding genes for bZip, HLH, HSF, GATA and Homeobox families. TSS was extended 1 kb upstream and downstream and split into 25 equally sized bins. (F) Proportion of peaks in *R. microsporus* genomic features. The same TF families as in (E) were considered.

A total of 50597 peaks were identified after filtering, with the number of peaks per TF ranging from 20 to several thousand (**Supplementary** Figure 3). Subsequently, we determined the motif bound by each TF, observing a high degree of similarity between experimental replicates (**Supplementary** Figure 4), even from those with FRIP > 5% in one replicate, supporting the reliability of the cistrome characterization. Clustering the identified motifs based on their similarity revealed that motifs from all HLH, GATA, Homeobox, HSF, and bZip were grouped together with TFs of the same family, indicating well-conserved target sequences within these families (**Figure 2C**).

Currently, only 307 motifs from fungal species are deposited in the public repository JASPAR^47^, with 292 derived from *Saccharomyces cerevisiae* and none from any EDF representative^47^. We queried our motif catalog against the JASPAR curated database and retained the closest matches for each of the 58 TFs based on motif similarity.

Sequences recognized by bZip TFs displayed some similarity to motifs bound by *S. cerevisiae* CRE (cAMP response element) factors Cst6/Aca2 and Sko1 (**Figure 2D**)^48–50^. HLH TFs bound to similar motifs to those recognized by Tye7 and Cbf1 (**Figure 2D**), which are also HLH TFs in *S. cerevisiae*^51,52^. Motifs from homeobox KN TFs also showed moderate similarity to the motif bound by *S. cerevisiae* Tos8, a TF involved in meiosis and cell damage response^53^. Interestingly, motifs bound by HSF TFs also displayed some similarity to those recognized by Mga1 and Hsf1 (**Figure 2D**), which are heat-shock factors involved in pseudohyphal growth and the essential regulation of heat-response genes in S. *cerevisiae*^54–56^. The observed similarities in this comparative analysis underscore the robustness of our dataset. In addition to identifying motifs, we analyzed TF binding relative to genomic features. Peaks tended to accumulate 0 to 250 bp upstream the TSS of protein-coding genes, with a large proportion located in promoter regions or the first exon (**Figures 2E** and **2F**). Noteworthy, HSF TF peaks accumulated closer to and downstream of the TSS (**Figure 2E**), suggesting distinct regulatory mechanisms among TF families. A genome-wide coverage analysis also revealed differences between TF families. While all GATA TFs were bound to similar regions (computing the sequencing coverage in 20 kb bins) notable differences were observed between the HSF and HLH TFs (**Supplementary** Figure 4), highlighting the diversity between all these TF families.

### The different TF families show target gene and functional diversification

To gain deeper insight into the role of each TF in gene expression regulation, we identified their putative targets. The 58 analyzed TFs displayed a wide range in the number of targets, from just six in the case of CP2-TF1 to several hundred for GATA Znf6, HOX2, or HLH Myc1 (**Supplementary Dataset 2**). Using this information, we generated a gene network including the 58 TFs studied and their respective predicted targets (**Figure 3A**). This network revealed well-differentiated groups of genes regulated by specific TF families. Notably, two distinct sets of genes regulated by GATA, HLH, Homeobox, and Fork-head TFs could be identified. In contrast, HSF and bZip TFs, with the sole exception of bZip5, were found to regulate a unique set of target genes (**Figure 3A**). Interestingly, the HMG-box TFs showed a remarkable heterogeneity in their target gene set, with a high degree of variability among the targets of the three TFs from this family that were analyzed (**Figure 3A**).

**Figure 3.**
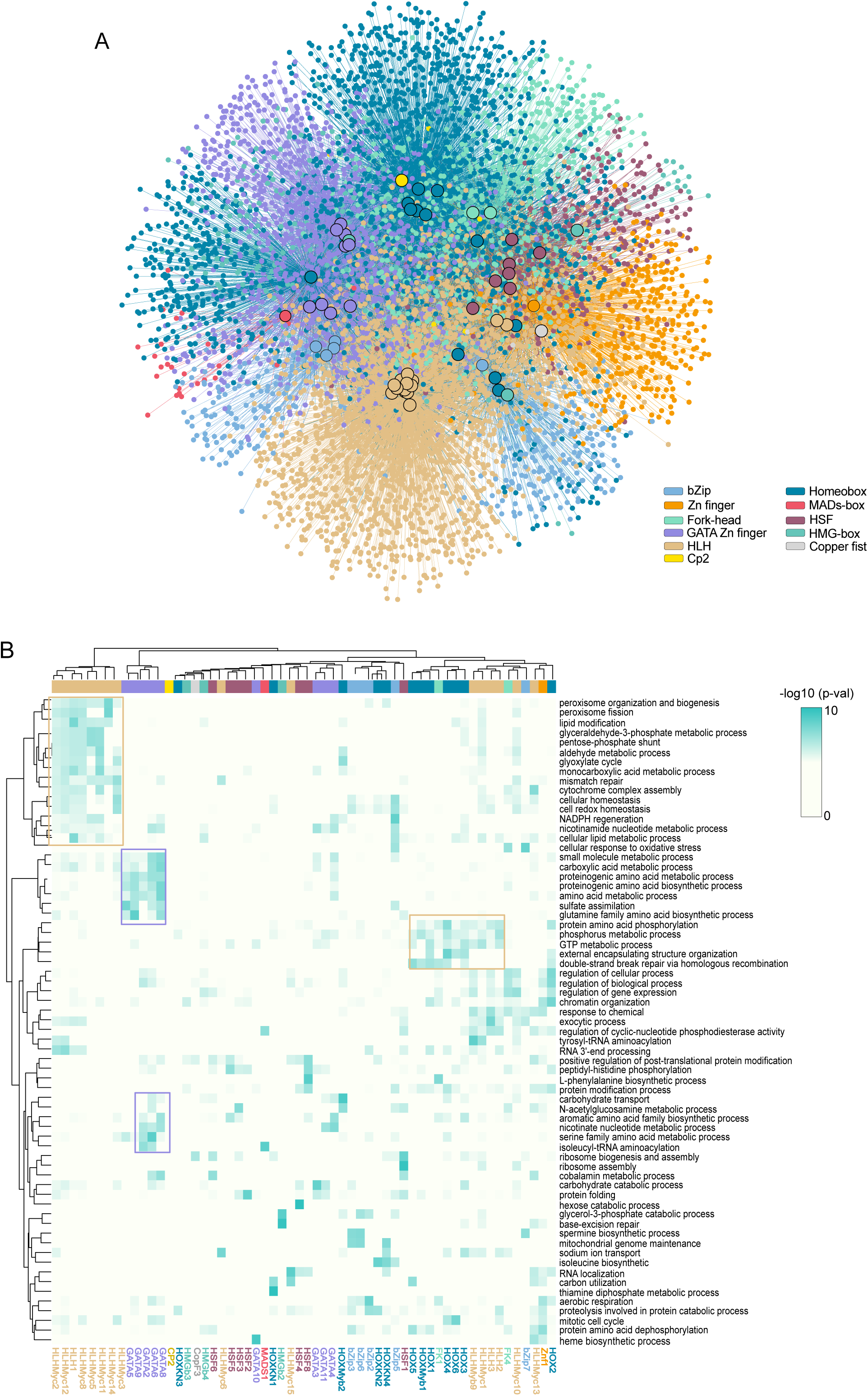
*R. microsporus* gene regulatory networks. (A) Gene network for the 58 TFs (bigger rounded circles) and their respective targets. TF families and and their targets are color-coded as indicated in the legend. (B) Functional classification of TF targets. The top 2 most enriched GO Biological Process terms for each TF were considered, and the p-value enrichment (-log10) was plotted for each term and each TF target set. White indicates no significant association was found for that term (p-val >0.05).

Given that the function of most TFs of *R. microsporus* is unknown, we performed functional enrichment analysis on the target gene sets for each TF. The targets of HLH TFs showed two well-defined clustering patterns associated with diverse biological functions, including lipid metabolism, redox homeostasis, and GTP metabolism (**Figure 3B**, orange squares), with some biological functions shared with Homeobox TF targets. GATA TF targets also showed a clear functional enrichment pattern, regulating genes involved in amino acid biosynthesis and other relevant processes, such as carbohydrate transport and sulfate assimilation (**Figure 3B**, purple squares). Since the TFs studied in this work are largely uncharacterized, this analysis also provided insights into their potential biological roles by examining their targets. For example, the TF Copf3 was found to target genes involved in ribosome biogenesis, while HLH Myc6 appeared to play a role in sodium ion transport (**Figure 3B**).

### The interplay between 6mA and TFs in gene expression regulation

Symmetric DNA 6mA plays an essential role in *R. microsporus*, with 35% of its genes containing MACs, including the 58 TFs identified in this study, and thus being directly regulated by 6mA (Lax et al, under revision in Nature Communications). Analyzing the targets of these 58 TFs could provide insights into the indirect role of this epigenetic modification in gene expression regulation. Peaks from the 58 TFs predicted that 6494 genes (60.1 % of total genes) were regulated by these TFs, whereas the total number of methylated genes in the same conditions was 3825 (35.7% of total genes). Comparing these two datasets revealed 2222 genes that were both regulated by TFs and 6mA, accounting for more than one-third of TF-regulated genes and almost 60% of all methylated genes (**Figure 4A**). This resulted in a total of 8097 genes (75.6% of total genes) directly or indirectly regulated by 6mA. Subsequently, we analyzed the targets of each TF in relation to the presence or absence of 6mA and found both individual and family-specific differences. The targets of HLH TFs, FK1, and Znf1 TFs were more frequently methylated than those of other TFs (**Figure 4B**). Further analysis of the binding signal of each TF across all MACs in the *R. microsporus* genome revealed a strong preference for binding to these regions by HLH Myc1, HLH Myc3, HLH Myc8, HLH Myc12, Znf1, and FK1 TFs (**Figure 4C**), which explains the aforementioned results.

**Figure 4.**
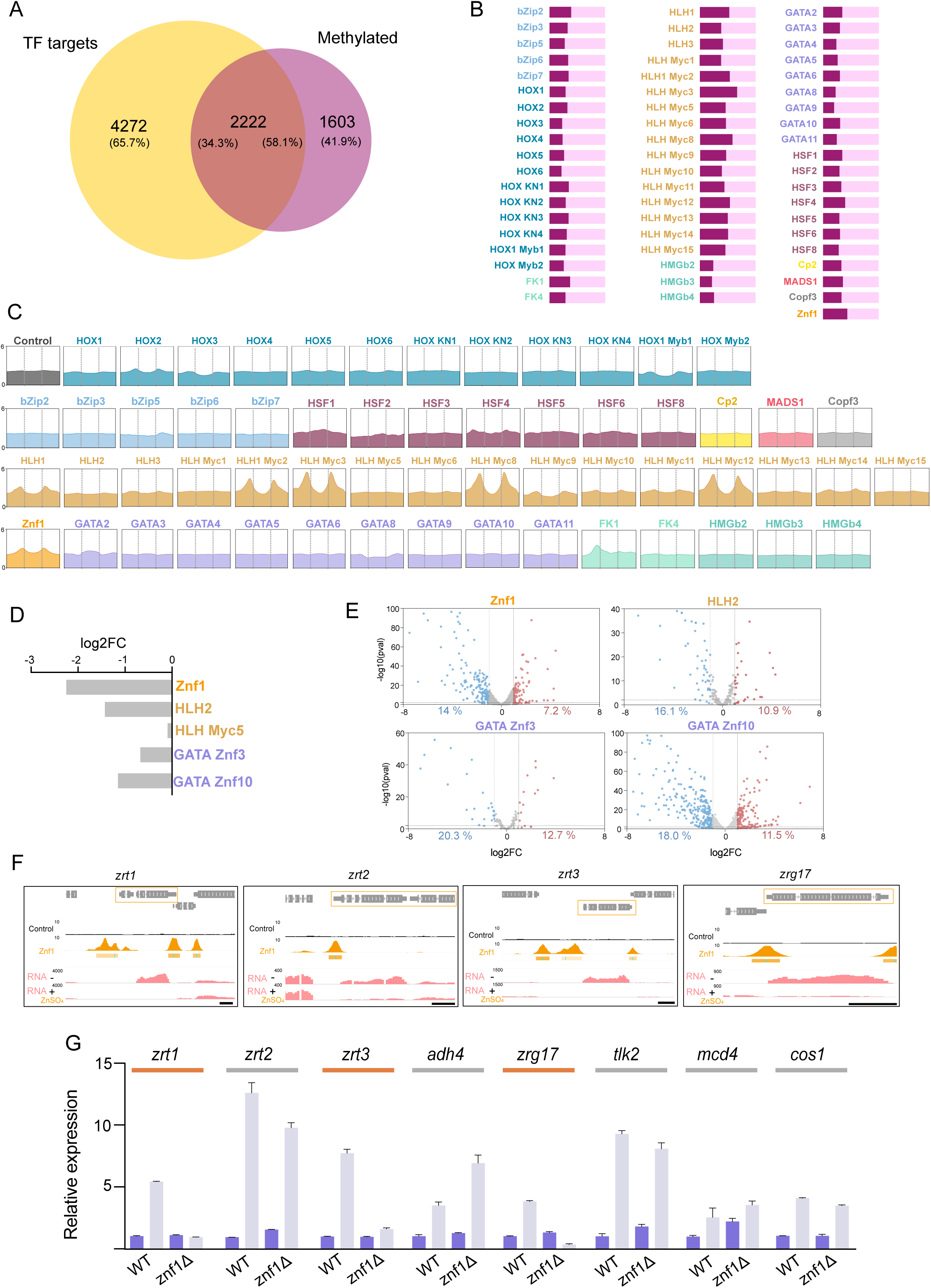
Indirect role of 6mA in gene expression regulation. (A) Venn diagram indicating TF target genes and methylated (6mA genes) in the *R. microsporus* genome. (B) Proportion of methylated (garnet) and unmethylated (light pink) genes for each TF-target set. (C) TF profile coverage over MACs. All MACs were sized to 500bp and extended 500 bp upstream and downstream. An empty backbone plasmid was used as a control (no protein expressed). Vertical dotted lines indicate MAC start and end. (D) Differential expression (log2FC) of genes demethylated when Mta1 expression is reduced. (E) Volcano plots for target genes of Znf1, HLH2, GATA Znf3 and GATA Znf10 TFs. log2FC and p-values are indicated for each gene. Only genes with log2FC |1 | and p-val < 0.05 were considered differentially expressed. (F) Genome view of genes whose promoter is bound by *R. microsporus* Znf1 and their expression is regulated by zinc. Scale bar = 500 bp. (G) Relative expression (qRT-PCR) of zinc-regulated genes for the wild-type and Znf1 mutants growing in zinc-rich media (purple) and zinc-depleted media (gray). Significant induction in the Znf1 mutant in the absence of zinc was tested using one-way ANOVA (p-val cut-off < 0.05). Those not induced were predicted as Znf1 targets and are indicated with an orange line.**Supplementary** Figure 5. (A) Profile coverage of all zinc finger TFs analyzed in this study over *zrt1* promoter. Coverage was plotted 1kb upstream of the *zrt1* TSS. An empty backbone plasmid was used as a control (no protein expressed). (B) Experimentally tested and likely *S. cerevisiae* Zap1 targets. The ID of the *R. microsporus* ortholog is indicated, as well as the log2FC, the genomic location, and the presence of Znf1 peak in their promoter (C) Schematic representation and results of PCR validation of the Znf1 mutant generation. Primers used to check the integration and homokaryosis are indicated as arrows in the scheme. (D) Growth of wild-type and Znf1 mutant in zinc-rich media (purple) and zinc-depleted media (gray). Image of tubes with 10 mL media is shown above the graph. A two-tailed Welch’s test was performed for each comparison indicated (ns = not significant). (E) Spore production quantification wild-type and Znf1 mutant in zinc-rich media (purple) and zinc-depleted media (gray). A two-tailed Welch’s test was performed for each comparison indicated (ns = not significant).

Given that all genes encoding the selected TFs were methylated, we investigated which lost methylation when Mta1 expression, the main methyltransferase involved in 6mA deposition in this fungus, was reduced (Lax et al. 2025, under revision in Nature Communications). Five of the studied TFs lost methylation (Znf1, HLH2, HLH Myc5, GATA Znf3 and GATA Znf10), leading to downregulation in four of them (**Figure 4D**). A total of 20 to 33% of their target genes were also differentially expressed (**Supplementary Table 2**). Moreover, we analyzed the effects on their respective target genes and found a significantly higher proportion of downregulated genes among the differentially expressed (DE) target genes of Znf1 and GATA Znf10 (**Supplementary Table 2** and **Figure 4E**), indicating that the loss of methylation also has an indirect effect on the targets of transcription factors regulated by this epigenetic modification.

### DAP-seq data provides insights into the regulatory mechanisms underlying important biological processes in EDF

The large number of analyzed TFs and the high degree of representativity of our dataset allowed us to explore the regulatory mechanisms underlying specific processes in *R. microsporus*, beyond the 6mA epigenetic regulation. As a proof of concept, we investigated the regulation of gene expression by zinc. Recently, we developed an expression system to control gene expression in Mucorales based on the use of the promoter of the *zrt1* gene, which encodes a zinc permease whose expression is dependent on zinc availability (Lax et al. 2025, under revision in Nature Communications). However, the TFs that regulate the expression of this and other genes related to zinc metabolism in EDF were unknown. Following the same approach previously used to analyze MACs, we scanned the promoter of the *zrt1* gene using the coverage information from all zinc finger TFs analyzed in this study, which could be potential regulators of this gene^57^ . We found a binding peak that the TF Znf1 upstream of the *zrt1* TSS (**Supplementary** Figure 5A). In *S. cerevisiae*, the ortholog of the *zrt1* gene is regulated by Zap1, a TF of C2H2/CCHC/CH/C5HC2 zinc finger family^58^. Znf1 is the only representative of this TF family in our dataset (**Supplementary Dataset 1**). Interestingly, we observed that several of the *R. microsporus* orthologs of Zap1-regulated genes involved in zinc metabolism^59,60^ (**Supplementary** Figure 5B) showed strong Znf1 binding peaks in their promoters and were regulated by zinc availability in the medium (**Figure 4F and Supplementary** Figure 5B), suggesting that they are Znf1 targets. To further investigate the role of *znf1* in *R. microsporus*, we generated a knockout mutant (**Supplementary** Figure 5C). Expression analysis by RT-qPCR of eight potential Znf1 targets in both the wild-type strain and the *znf1* mutant under zinc-depleted and zinc-replete conditions revealed that the *zrt1*, *zrt3*, and *zrg17* orthologs were not induced in the mutant under zinc-depleted conditions, suggesting a direct regulatory role of Znf1 in their expression (**Figure 4G**). The remaining potential Znf1 targets displayed expression patterns similar to those in the wild-type strain, suggesting functional redundancy with other TFs contributing to their regulation in *R. microsporus* (**Figure 4G**). Disruption of *znf1* resulted in reduced growth under zinc-depleted conditions compared to the wild-type strain (**Supplementary** Figure 4D), while the sporulation rate remained unaffected (**Supplementary** Figure 4E).

Another aspect that strongly captured our interest during the analysis of the *R. microsporus* cistrome was the regulation of sexual reproduction. Zygomycete fungi are characterized by having two distinct mating types, distinguished by the presence of the *sexP* (+) or *sexM* (-) genes in their mating locus. These genes encode HMG-box TFs that regulate mating-specific genes in each type^61,62^. While some target genes of *sexP* and *sexM* are known^61^, the TFs that bind to their promoters and regulate their expression remain unidentified. Using the data from all the TFs analyzed, we scanned the promoter region of the *sexP* gene in *R. microsporus* and found a robust binding peak for bZip5 (**Supplementary** Figure 6). This TF, which binds to regions and regulates genes different from the other bZip factors analyzed (**Figure 3A and Supplementary** Figure 4), presents a promising candidate for studying gene regulation in the sexual reproduction of these fungi. Unfortunately, the *R. microsporus* strain used in this study (ATCC 11559) has not successfully undergone mating under laboratory conditions, so further investigations using other strains or species will be necessary to elucidate the involvement of bZip5 in the regulation of reproduction.

### DAP-seq data unveils white-collar control of light-regulated genes in *R. microsporus*

The set of TFs analyzed by DAP-seq included a White Collar-2 (WC-2) protein of the WC-2C/D type, here referred to as GATA Znf10 or WC-2C (**Supplementary** Figure 7A), making possible the identification of blue light-responsive genes regulated by the white-collar photoreceptor composed of WC-2C and an uncharacterized white collar-1 protein^63^ . While the genetic regulation of light responses has been extensively studied in Mucorales^64–69^ and the biological role of white-collar proteins is well-documented^63,70,71^, the genes directly or indirectly regulated by these photoreceptors in this fungal group remain unknown. We detected strong binding signals of WC-2C to 1,394 potential target genes (**Supplementary Dataset 2**), with particular enrichment in the promoter region (**Figure 5A**). The binding motif was identified as GATC (**Figure 5B**), which is consistent with the consensus binding sequence described for white collar proteins in fungi^21,72,73^.

**Figure 5.**
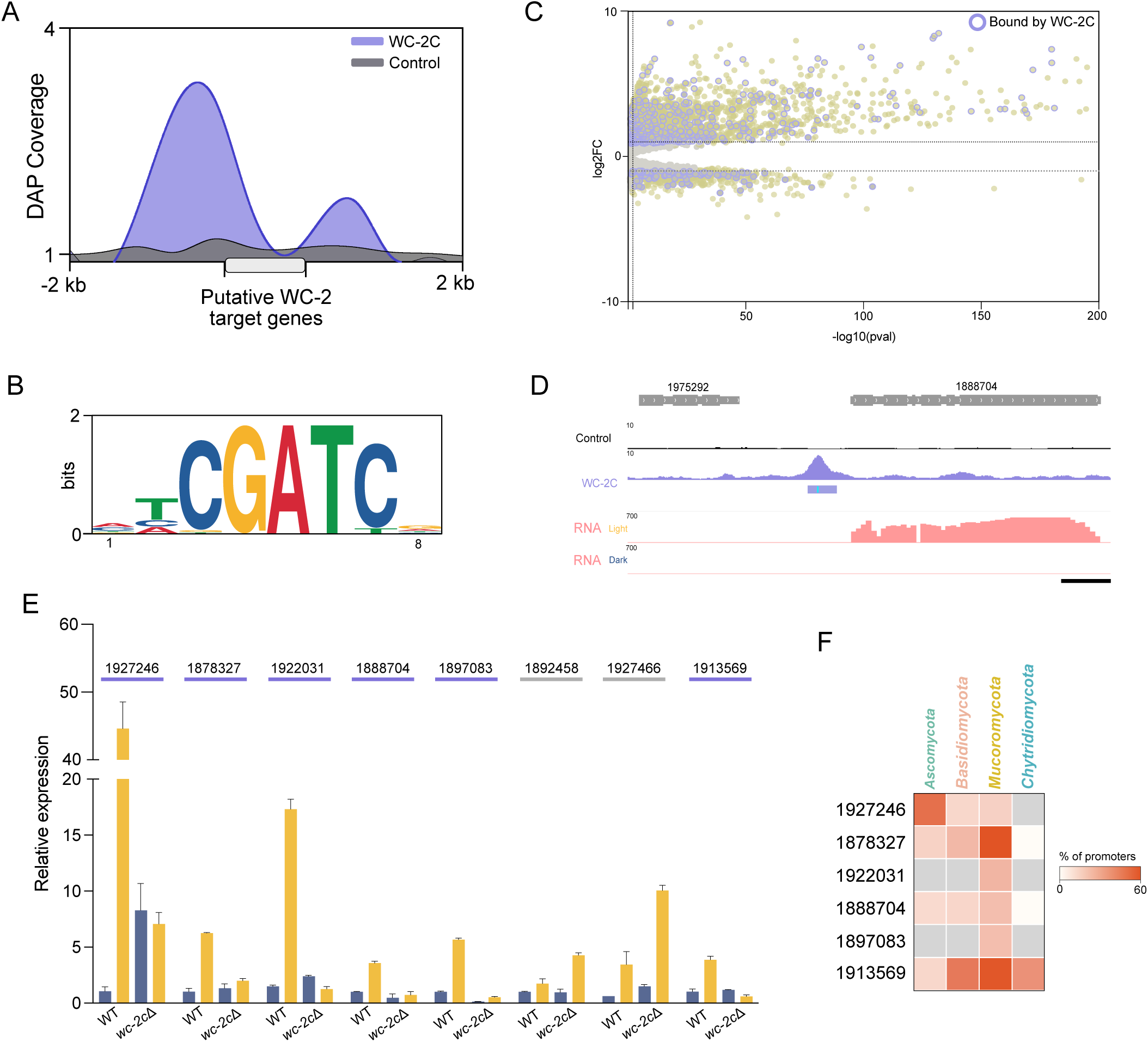
Wc2 implication in light-regulated genes. (A) WC-2C (GATA Znf10) and control (empty backbone plasmid) DAP coverage over predicted WC-2C targets. Genes were sized to 1 kb and coverage was computed over genes and 2 kb upstream and downstream. (B) Motif bound by WC-2C in *R. microsporus* obtained using the 100 most significant peaks, as scored by signal value in column 7 of the narrowPeak file from WC- 2C. (C) Ligh-induced and repressed genes regulated by WC-2C. A volcano plot shows all differentially expressed genes (log2FC |1 | and p-val > 0.05) in yellow and those with a WC-2C peak in their promoter rounded with a purple circle. (D) Genomic snapshot showing the WC-2C coverage and RNA-seq expression in light and dark growth conditions for the 1888704 gene. Scale bar = 500 bp. (E) Relative expression of zinc- regulated genes (qRT-PCR) for the wild-type and *wc-2c* mutants growing in the dark (dark blue) and under light exposure (yellow). Significant induction in the WC-2C mutant in the presence of light was tested using one-way ANOVA (p-val cut-off < 0.05). Those not induced were predicted as WC-2C targets and are indicated with a purple line. (F) Percentage of orthologous promoters with the WC-2C motifs. Promoter (600 bp) was extracted from each ortholog and scanned with FIMO using the *R. microsporus* WC-2C motif.

To identify the light-responsive genes regulated by the blue-light photoreceptor containing WC-2C, we compared its predicted targets with the genes regulated by light in *R. microsporus* (Lax et al. 2025, under revision in Nature Communications). Among the 2,871 genes regulated by light, 427 genes (14.8%) showed a WC-2C binding peak in their promoter regions. Of these, the majority—344 genes—were induced, while 83 genes were repressed (**Figure 5C**), indicating that WC-2C predominantly functions as a transcriptional activator in this fungus. To validate the involvement of WC-2C in regulating these genes, we generated a mutant strain with a disrupted *wc-2c* gene (**Supplementary** Figure 7B) and analyzed the expression of eight light-responsive genes in both the wild-type and mutant strains under light and dark conditions. Six of these genes failed to be upregulated in response to light in the mutant (**Figure 5E**), confirming that WC-2C is critical for the light-responsive expression of most genes it binds. Notably, the *wc-2c* mutation resulted in increased sporulation under dark growth conditions compared to the wild-type strain (**Supplementary** Figures 7C and 7D). In addition, the mutant exhibited reduced aerial mycelium development and an increased accumulation of sporangiophores at the base of the mycelium (**Supplementary** Figure 7E). This phenotype is consistent with the loss-of-function of the light-inducible *crgA* gene (ID 1913569) in *R. microsporus*^74^, one of the WC-2C targets that failed to increase expression in the *wc-2c* knockout mutant (**Figure 5E**).

One of the most interesting insights provided by the DAP-seq data is the opportunity to study gene regulation and evolutionary conservation of the targets. Having identified six direct WC-2C targets in this study, we analyzed whether white-collar proteins also regulate the orthologs of these genes in other fungi. To do this, we identified orthologous genes of each target in the 303 fungal genomes available in FungiDB^75^ and searched for the GATC motif in their promoters. In addition to Mucoromycota, we detected a high frequency of the WC-2C-recognized motif in the Ascomycota and Basidiomycota, especially for genes ID: 1927246, 1878327, and 1888704, suggesting a possible regulation of these genes by WC-2C in other fungal species (**Figure 5F**). Moreover, we found a high frequency of the GATC site in the promoters of *crgA* (ID 1913569) orthologs, including those from Chytridiomycota (**Figure 5F**), indicating strong conservation of the *crgA* regulation by WC-2C across fungal clades.

### Exploring evolutionary conservation of promoter architecture across fungal lineages

The conservation of targets observed for WC-2C and *crgA* prompted us to examine the conservation of the promoter structure across distinct fungal clades more closely. We examined WC-2C motif occurrence and found that similar to the peak detected in *R. microsporus*, other Mucoromycota representatives also contained the GATC motif located 300 bp upstream the TSS (**Figure 6A**). Interestingly, the specific distribution of this motif was also shared by some Basidiomycota and Ascomycota species, such as *Kwoniella dejecticola* and *Aspergillus carbonarius*, suggesting a certain degree of conservation in promoter architecture. In contrast, distinct promoter architecture signatures were observed for each clade. Ascomycota displayed two GATC sites located ∼150 bp upstream of the TSS, separated by 20-30 bp (**Figure 6A**), consistent with previous characterizations of WC-2 binding to *Neurospora* promoters^32^. In basidiomycete orthologs, GATC sites were located around 100-150 bp upstream of the TSS, with closer spacing compared to those detected in Ascomycota (**Figure 6A**). In the chytrid *Batrachochytrium dendrobatidis,* only one GATC site was detected, coinciding with site distribution in Basidiomycota.

**Figure 6.**
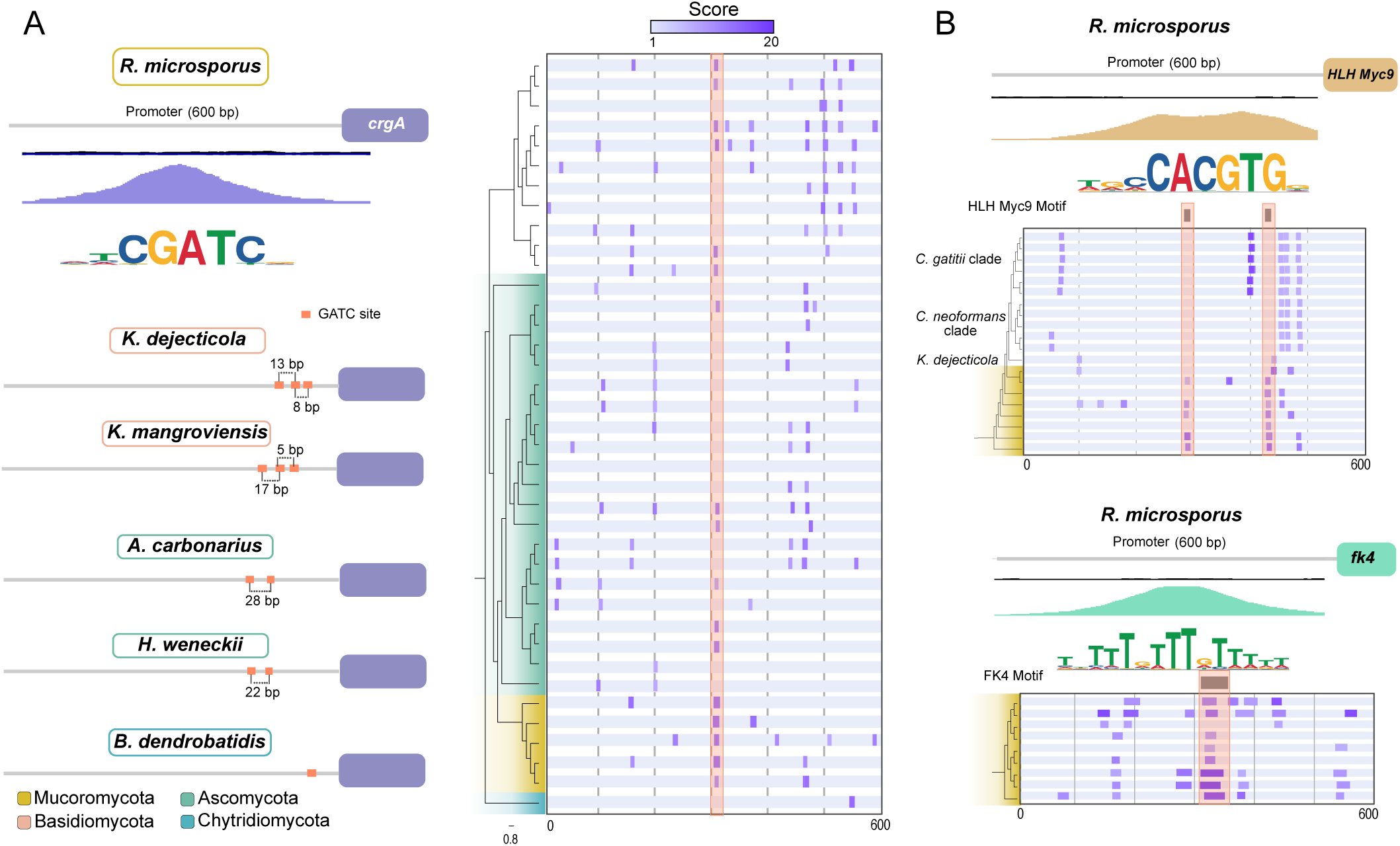
Promoter structure evolution in fungi. (A) Analysis of the *crgA* promoter across the fungal kingdom. A schematic diagram is shown on the left for *R. microsporus* on the top and representative species for each phylum. Distribution of WC-2 motifs for each promoter is indicated on the right panel. The orange line indicated the distribution of the GATC site conserved across all Mucoromycota and bound by WC-2 in *R. microsporus* according to DAP-seq data. Sites are color-coded according to the FIMO score. (B) HLH Myc9 (top) and FK4 (bottom) motifs distribution on its own promoters. The orange line indicated the distribution of the HLH Myc9 and FK4 sites conserved in Mucoromycota and bound by these TFs in *R. microsporus* according to DAP-seq data. Sites are color-coded according to the FIMO score. Information on the genomes analyzed can be found in Supplementary Dataset 1.

Transcriptional autoregulation is one of the simplest yet most efficient mechanisms of gene expression regulation. Autoregulation loops are highly conserved throughout vertebrate evolution^76^ and are present from the lambda phage to higher complex eukaryotes^77^. We scanned the promoters of each TF studied in this work for signs of autoregulation (**Supplementary** Figures 8A and 8B). To investigate the conservation of these autoregulation mechanisms and characterize promoter architecture conservation, we focused on two TFs, HLH Myc9 and FK4, which showed evidence of autoregulation. The promoter of the HLH Myc9 TF displayed a robust peak with two binding sites (**Supplementary** Figure 8A **and Figure 6B**). The promoter HLH Myc9 ortholog in Mucoromycota also contained a binding site located near the TSS, although the presence of a second motif was more variable (**Figure 6B**). Notably, this second site was also missing in the basidiomycete *Kwoniella dejecticola,* though the site located ∼100 bp upstream of the TSS was conserved (**Figure 6B**). In contrast, the promoters of HLH Myc9 orthologs in *Cryptococcus* species showed a completely different architecture, with the *C. gattii* clade displaying a site located 200 bp upstream of the TSS that was absent in the C*. neoformans* clade, indicating inter-species variability in promoter architecture, which could also suggest functional divergencies in the TF orthologs. Orthologs of FK4 were only found in Mucoromycota fungi, and all their promoters contained the motif bound by FK4 (**Figure 6B**), suggesting that the autoregulation loop of this TF is well-conserved within this early diverging fungal phylum.

## Discussion

The comprehensive characterization of cistromes has long been hindered by technical challenges and the limited scalability of procedures for identifying TFDS^8^. However, the recent development of the DAP-seq technique has significantly increased the scalability and reliability of fungal cistrome characterizations, democratizing its application to groups of organisms that have historically been understudied^13,15,31,78,79^. A notable example is the EDF, a fungal subkingdom that has traditionally received less attention than Dikarya, a disparity largely due to difficulties in genetic manipulation^80^ and the complexity of laboratory handling^34^. Despite these challenges, the limited knowledge of their biology makes EDF particularly valuable for understanding the evolution of eukaryotic organisms. Coupled with their immense diversity^35^, EDF provide ideal models for exploring new regulatory mechanisms. In this context, the characterization of 58 TFs from *R. microsporus* represents a major step forward in understanding regulatory mechanisms in the early branches of fungal evolution.

The TF landscape in fungi shows remarkable diversity both between EDF and Dikarya and within the phyla included in EDF. This diversification highlights the evolutionary dynamics of TFs across fungal lineages, offering new insights into their regulatory complexity. The generation of genome-wide binding profiles for 58 TF belonging to 11 different families using DAP-seq has led to the identification of the DNA motifs recognized by these TFs, significantly expanding the catalog of fungal TF motifs, which has been dominated by motifs from Dikarya^47^. Analysis of the binding profiles and peak distribution across the genome revealed interesting differences between TF families, suggesting the presence of specific regulatory mechanisms, a phenomenon observed in higher organisms^81^ . Moreover, the cistrome characterization of *R. microsporus* has further elucidated the complex interactions between TFs and their genomic targets (**Figure 3A**). Functional classification of these targets revealed a robust clustering of TFs belonging to the same family (**Figure 3B**), highlighting the existence of functional specialization among TF families.

6mA is an essential epigenetic modification in *R. microsporus* and is associated with transcriptionally active genes (Lax et al. 2025, under revision in Nature Communications). In recent years, the biological implications of this epigenetic modification have garnered significant interest, with studies spanning from humans and mice to plants and fungi^42–44,82–86^. Although approximately 30% of the genes in *R. microsporus* are directly regulated by 6mA (Lax et al. 2025, under revision in Nature Communications), the indirect effects of this modification, particularly through the methylation of genes encoding TFs and the regulatory networks that they control, have remained largely unexplored. In this study, we show that 58 TFs collectively target 4272, representing 40% of the *R. microsporus* genome. This finding suggests that 6mA exerts indirect control over a substantial portion of the genome through its regulation of TFs. The essential nature of 6mA complicates efforts to analyze how this epigenetic modification affects the expression of TFs it regulates. However, a conditional mutant for the catalytic subunit of the 6mA methyl transferase complex shows a ∼30 to 40% reduction in 6mA under restrictive conditions (Lax et al. 2025, under revision in Nature Communications). When we examined the methylation of genes encoding the 58 TFs studied, a subset, likely the most sensitive to the loss of methyltransferase activity, showed reduced methylation and decreased expression. Interestingly, the expression of a significant number of target genes regulated by these TFs was altered, with downregulation observed more frequently. These results provide strong evidence that 6mA indirectly controls target genes through its regulation of TFs, highlighting its pivotal role in the transcriptional regulatory networks of *R. microsporus*.

The regulatory influence of 6mA extends even further when considering both direct targets and indirect targets, encompassing nearly three-quarters of the genes in the *R. microsporus* genome. This extensive control likely explains why the loss of symmetric 6mA in this fungus is lethal. Additionally, a proportion of methylated genes are targeted by TFs regulated by 6mA. Interestingly, the methylation status of target genes varies significantly among different TFs and families, with HLH-type TFs showing the strongest preference for binding to methylated genes. This differential affinity for methylated versus unmethylated genes has already been demonstrated in a large-scale study using DAP-seq to investigate genes with and without 5mC in *A. thaliana*^13^.

One of the interesting aspects of the DAP-seq data generated in this study is its ability to evaluate any genomic region for binding by a wide range of TFs. By using this information, we expand our understanding of biologically relevant processes that have been poorly or scarcely characterized in EDF. Although several light-responsive genes have been characterized in Mucorales^64,65,87^ and changes in gene expression associated with loss-of-function mutations in white-collar proteins have been described^66,70,71,88^, it has been the integration of gene expression data from light and dark growth conditions (Lax et al. 2025, under revision in Nature Communications) with DAP-seq data for WC- 2C, one of the TFs analyzed in this study, that has enabled the identification and validation of several target genes. This approach provides a better understanding of the role of this photoreceptor subunit in the transcriptional response to light, including its regulation of the carotenogenic repressor *crgA*. Similarly, while zinc-regulated systems have been exploited to develop regulatable expression systems in *R. microsporus* (Lax et al. 2025, under revision in Nature Communications), the TFs involved in this regulation remained largely unknown. The characterization of the Znf1, a Zap1-like TF involved in zinc- related transcriptional regulation in yeasts^59^, reveals its role in regulating several genes involved in zinc metabolism in *R. microsporus*. However, the orthologs of some Zap1 targets are not regulated by Znf1 in *R. microsporus*, highlighting differences in regulatory mechanisms and suggesting the involvement of other, yet unidentified, transcription factors in the regulation of these genes. On another note, despite the significant peak found for bZip5 in the promoter of the *sexP* gene, further studies are required to confirm the involvement of this TF in regulating the mating locus, particularly using *R. microsporus* strains with known mating conditions and compatible partners are available^89^.

The identification of TFBS provides valuable information, not only for uncovering additional targets within the same organism but also for predicting regulatory mechanisms in other species and exploring the conservation of promoter structures across the fungal kingdom. By focusing on the high conservation of autoregulation loops^76,77^, we have identified both common regulatory elements and specific signatures across different fungal phyla. Overall, these results highlight the broad relevance of the data generated, not only for advancing our understanding of *R. microsporus* biology but also for enriching our knowledge of gene regulation mechanisms across the fungal kingdom.

## Methods

### DAP-seq

DNA Affinity Purification sequencing (DAP-seq) experiments were performed following a previously established procedure^13,14^ with minor modifications^15^. Briefly, genomic DNA from *R. microsporus* was fragmented to an average size of 150bp (Covaris LE220- Plus) and sequencing libraries were prepared using the KAPA HyperPrep kit (Roche). In parallel, genes encoding transcription factors were codon refactored to overcome synthesis constraints (**Supplementary Dataset 3**) and assembled from synthetic DNA fragments (Twist Biosciences, CA), cloned into pIX_HALO plasmid backbone yielding an in-frame N-terminus HALO tag, confirmed via sequencing, and used for PCR amplification to yield linear PCR fragments containing the Halo-TF fusion protein driven by a T7 promoter. The PCR product was purified with SPRI beads, and the correct size of each was verified using a Fragment Analyzer (Agilent Technologies). For each DAP-seq reaction, each expressed protein was used, along with 20 ng of the prepared fragment library and 2000 ng of salmon sperm DNA to mitigate non-specific binding. An empty backbone plasmid was used as a control. The resultant DAP-seq libraries were pooled for sequencing on a NovaSeq using the S4 flowcell (Illumina).

### DAP-seq data processing

DAP-seq reads were mapped to the *R. microsporus* reference genome (https://mycocosm.jgi.doe.gov/Rhimi59_2/) using Bowtie2^90^. MACS3^91^ (v3.0.0a6) was used to call the peaks and motifs were generated using MEME^92^ with the 100 most significant peaks, as scored by signal value in column 7 of the narrowPeak files. TFs were filtered, discarding those with FRIP (Fraction of Reads in Peaks) < 5%. The top motif for each TF was clustered and visualized using the R package MotifStack (v1.48.0)^93^. Peak distribution was assessed using Bedtools (v2.30.0)^94^ and visualized with deepTools (v3.1)^95^. Whole-genome genome comparisons and binding profiles over regions of interest were performed using deepTools (v3.1)^95^ using a 10 kb bin size for whole-genome coverage comparisons and 10 bp for binding plots. TF targets were predicted using the RnaChipIntegrator tool^96^, filtering only those with a peak located at -500 + 100 bp from TSS. The gene network of TF and TF targets was constructed using Cytoscape (v3.10.2)^97^. GO annotations were retrieved from the *R. microsporus* v2 annotation available at Mycocosm (https://mycocosm.jgi.doe.gov/Rhimi59_2). A p-value of 0.05 was used as a cutoff for the identification of enriched terms (Fisher’s exact test). The top 2 most enriched GO Biological Process terms were considered for each TF-target set. Motifs were compared against fungal motifs in the JASPAR database using TomTom (MEME suite v 5.5.5)^92^. The best hit for each TF was retained and the p-value of the significance was used for plotting. Promoters were scanned using FIMO (MEME suite v 5.5.5)^92^, and only motifs with scores > 0 were retained. Scores were used to color code the motifs found.

### Phylogenetic and protein conservation analyses

InterProScan^98^ domain data was retrieved from sequenced and annotated genomes available at JGI (https://mycocosm.jgi.doe.gov/mycocosm). The complete list of TF families and species included in the analysis is indicated in Supplementary Data 1. The species tree was generated after analyzing the 62 proteomes with OrthoFinder (Inflation factor, -I 1.5)^99^. The white-collar phylogenetic tree was constructed aligning known *Mucor lusitanicus* and *Phycomyces blakesleeanus* white-collar proteins^63^ with the predicted *R. microsporus* WC-2 using MAFFT^100^. The phylogenetic tree was obtained using RAxML^101^ (PROTGAMMAWAGF substitution model) with 100 bootstrap replicates.

### qRT-PCR

The expression of the potential TF target genes was analyzed by qRT-PCR at different Zinc concentrations (ZnSO4) (0 mg/L and 20 mg/L) and in dark growth and with light exposure (24 hours). Amplicons were generated with specific primers (Supplementary Table 3). Total RNA was extracted using the RNeasy Mini kit (Qiagen) and submitted to a DNase treatment (Sigma, On-Column DNaseI treatment set). cDNA was synthesized from 1 μg of total RNA using the iScript^TM^ cDNA synthesis kit (Biorad). cDNA amplification was carried out in triplicate using a reaction mixture containing 2x Power SYBR® Green Master Mix (Applied biosystem, Waltham, MS, USA), 150 nM of gene-specific primers, and 100 ng of cDNA. The real-time PCR was carried out using the QuantStudio^TM^ 1 real-time PCR system (Applied Biosystems) according to the established experiment template in the equipment. The melting curve and non-template controls were also measured to discern non-specific amplifications. The relative expression of target genes was normalized with the amplification levels of the constitutively expressed *mitC* (Mitochondrial carrier) gene (ID:1927801) (Supplementary Table 3) and calculated using the 2−ΔΔCT method.

### Mutant strain generation and validation

*R. microsporus* mutant strains were generated following previously established procedures^74,102^. For *wc2* (ID: 1921537) and *znf1* (ID: 1799646) disruption, a template DNA containing the *pyrF* (ID: 1869696) gene and flanked by 38 nt homology regions was used to disrupt the targeted gene. Recombination was enabled through the directed double-strand break (DSB) provoked by Cas9 in an uracil-auxotrophic strain (UM1)^74^. Designed crRNAs (Alt-R^TM^ CRISPR-Cas9 crRNA) used to target the CRISPR/Cas9 system were purchased from IDT (Supplementary Table 4). crRNAs were coupled with tracRNA (Alt-R^TM^ CRISPR-Cas9 tracRNA, IDT) to form the gRNA, which was assembled in vitro with Cas9 (Alt-R^TM^ S.p. Cas9 nuclease, IDT) to form the ribonucleoprotein complex. Gene disruptions and homo/heterokaryosis was validated by PCR using primers flanking the disruption site that amplifies both WT and mutant nuclei if present. All primers used for construct amplification and PCR validation are listed in Supplementary Table 3. DNA was extracted following a phenol/chloroform extraction^103^, and all PCR amplifications were performed with the Herculase II Fusion DNA polymerase (Agilent), adapting the annealing temperature to each primer pair following the manufacturer’s recommendations.

### Fungal growth conditions

All the fungal strains used and generated in this work derive from *Rhizopus microsporus* ATCC 11559. Transformants of the auxotrophic strain UM1 strain^74^ using the *pyrF* marker were grown in Minimal Media with Casamino acids (MMC)^104^. During transformation, electroporated protoplasts were resuspended in ice-cold Yeast Peptone Glucose (YPG) media + 0.5 M Sorbitol for 90 min and then centrifugated at 800 rpm and resuspended in YNB media +0.5 M Sorbitol. All strains were grown at 30°C under light or dark conditions as specified. DAP-seq and gene expression samples were collected 24 hours after inoculating 1.10^6^ spores per plate. To analyze the expression of the zinc-regulated genes, the medium was supplemented with 20 mg/L of ZnSO4.

### Phenotypic characterization

Dark and light exposure comparisons were performed by growing 1.10^4^ spores distributed in five points of the plate in YPG medium pH 4.5. Plates were incubated at 30°C for 72 hours, exposed to continuous white light, or covered in foil. Plates were imaged using a Stemi 305 Zeiss microscope (integrated camera). For sporulation quantification, 1.10^6^ spores were inoculated in the center of a plate of YPG medium pH 4.5. Plates were incubated at 30°C for 72 hours. A chunk of agar of 1cm 2 (one per plate, three for each strain) was transferred to a 50 ml tube containing 10 mL of PBS 1X. The tube was vigorously vortexed to release spores. Spores were counted in a Neubauer chamber to determine the concentration, and total spore production was calculated.

For *znf1*Δ phenotypic characterization, 1.10^6^ spores were inoculated in a flask with media with and without ZnSO4 supplementation (20mg/L). Biomass was collected after 72 hours of growth in a rotary shaker at 30°C and lyophilized. Dry weight was obtained from three independent growth experiments. Spore production was performed as described above but with media supplemented with ZnSO4 when indicated.

### Statistical analysis

Statistical details are detailed in the Results, Figures, and Figure legends, including the number of biological and technical replicates as well as the dispersion and precision measures (mean and SD). Statistical analyses were performed using GraphPad Prism 9 (https://www.graphpad.com). Data normality was analyzed using the Shapiro-Wilk normality test with a significance level (alpha) of 0.05. Direct comparisons between the expression levels of methylated and unmethylated genes were analyzed using Welch’s test. Pearson correlation factors, p-values cut-off for Fisher exact tests, and Welch’s tests for direct comparisons are indicated in the respective figure legends. To assess the significance of downregulated genes of unmethylated TF target genes, contingency tables with upregulated and downregulated genes were generated for target and not target genes and Fisher exact test was conducted with a significance level (alpha) of 0.05 (two-tailed). Differences in TF proportions between species were evaluated by creating contingency tables and performing Chi-square comparisons. Two-tailed p values were calculated setting 0.05 as the threshold for significant differences.

## ACKNOWLEDGMENTS

This research was funded by the MCIN/AEI/ 10.13039/501100011033 by “ERDF A way of making Europe,” by the “European Union” grant PID2021-124674NB-I00 to F.E.N. and V.G. The work (10.46936/10.25585/60001127) was conducted by the US Department of Energy Joint Genome Institute (https://ror.org/04xm1d337), a DOE Office of Science User Facility, was supported by the Office of Science of the US Department of Energy under Contract No. DE-AC02-05CH11231.

## Author contributions

C.L. conducted most of the analyses, prepared the *R. microsporus* material for sequencing, performed bioinformatic analysis of the results, prepared the]igures and tables, designed and coordinated the project, and wrote the manuscript with signi]icant input from G.T., F.E.N, and V.G. L.A.B. participated in the processing of DAP-seq data. G.T. participated in qRT-PCR analyses and bioinformatic analyses. N.L.-M. participated in qRT-PCR analysis. Y.Z. performed DAP-seq. I.K.B. designed and synthesized transcription factors for the DAPSeq experiments. V.N. managed the project. E.N. managed the project and provided materials. F.E.N. supervised the study. V.G. analyzed the results and designed, supervised, and coordinated the project.

## Data and code availability

DAP-seq and RNA-seq data are available at Sequence Read Archive (SRA) under the accession number SRP496793, SRP496795, SRP496796, SRP496797, SRP496798, SRP496799, SRP496800, SRP496816, SRP496817, SRP496818, SRP496819, SRP496820, SRP496821, SRP496822, SRP496823, SRP496824, and SRP496825.

Custom scripts generated for data processing are available at https://github.com/ghizlanetahiri95/DAP_Rhizopus.

