## Supplementary information for "Genome-wide exploration of the transcriptional regulatory landscape in the early-diverging fungus *R. microsporus* reveals pervasive DNA methyl adenine regulatory network"

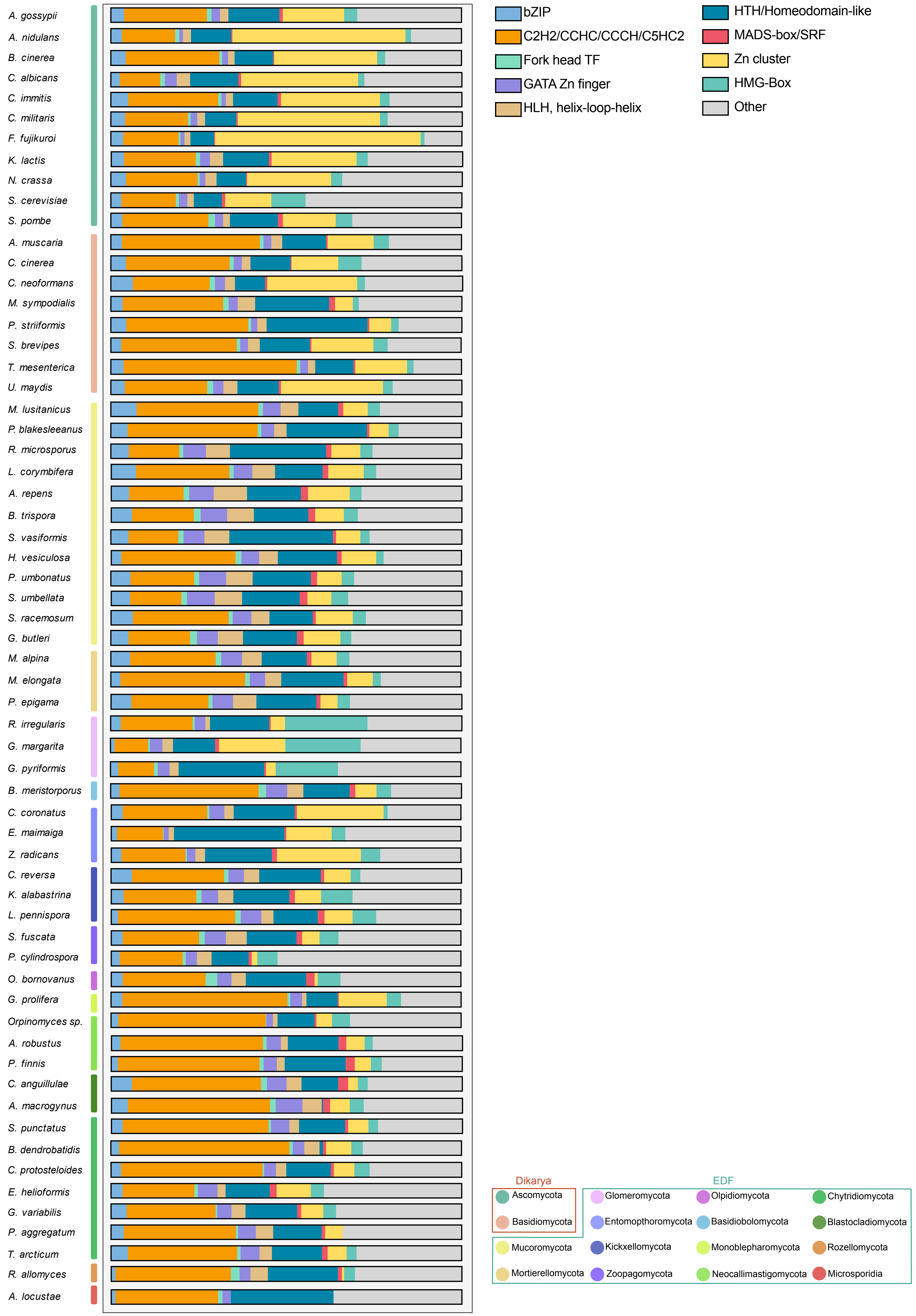
A. gossypii
A. nidulans
B. cinerea
C. albicans
C. immitis
C. militaris
F. fujikuroi
K. lactis
N. crassa
S. cerevisiae
S. pombe
A. muscaria
C. cinerea
C. neoformans
M. sympodialis
P. striiformis
S. brevipes
T. mesenterica
U. maydis
M. lusitanicus
P. blakesleeanus
R. microsporus
L. corymbifera
A. repens
B. trispora
S. vasiformis
H. vesiculosa
P. umbonatus
S. umbellata
S. racemosum
G. butleri
M. alpina
M. elongata
P. epigama
R. irregularis
G. margarita
G. pyriformis
B. meristorporus
C. coronatus
E. maimaiga
Z. radicans
C. reversa
K. alabastrina
L. pennispora
S. fuscata
P. cylindrospora
O. bornovanus
G. prolifera
Orpinomyces sp.
A. robustus
P. finnis
C. anguillulae
A. macrogynus
S. punctatus
B. dendrobatidis
C. protosteloides
E. helioformis
G. variabilis
P. aggregatum
T. arcticum
R. allomyces
A. locustae
bZIP
C2H2/CCHC/CCCH/C5HC2
Fork head TF
GATA Zn finger
HLH, helix-loop-helix
HTH/Homeodomain-like
MADS-box/SRF
Zn cluster
HMG-Box
Other
Dikarya
Ascomycota
Basidiomycota
Mucoromycota
Mortierellomycota
EDF
Glomeromycota
Entomopthoromycota
Kickxellomycota
Zoopagomycota
Olpidiomycota
Basidiobolomycota
Monoblepharomycota
Neocallimastigomycota
Chytridiomycota
Blastocladiomycota
Rozellomycota
Microsporidia

**Supplementary Figure 1.** Proportions (Total number of TF from a family/ Total number of TFs) of the main fungal TF families indicated for each species. The different phyla are color-coded as indicated in the legend. Information on the genomes analyzed can be found in Supplementary Dataset 1.

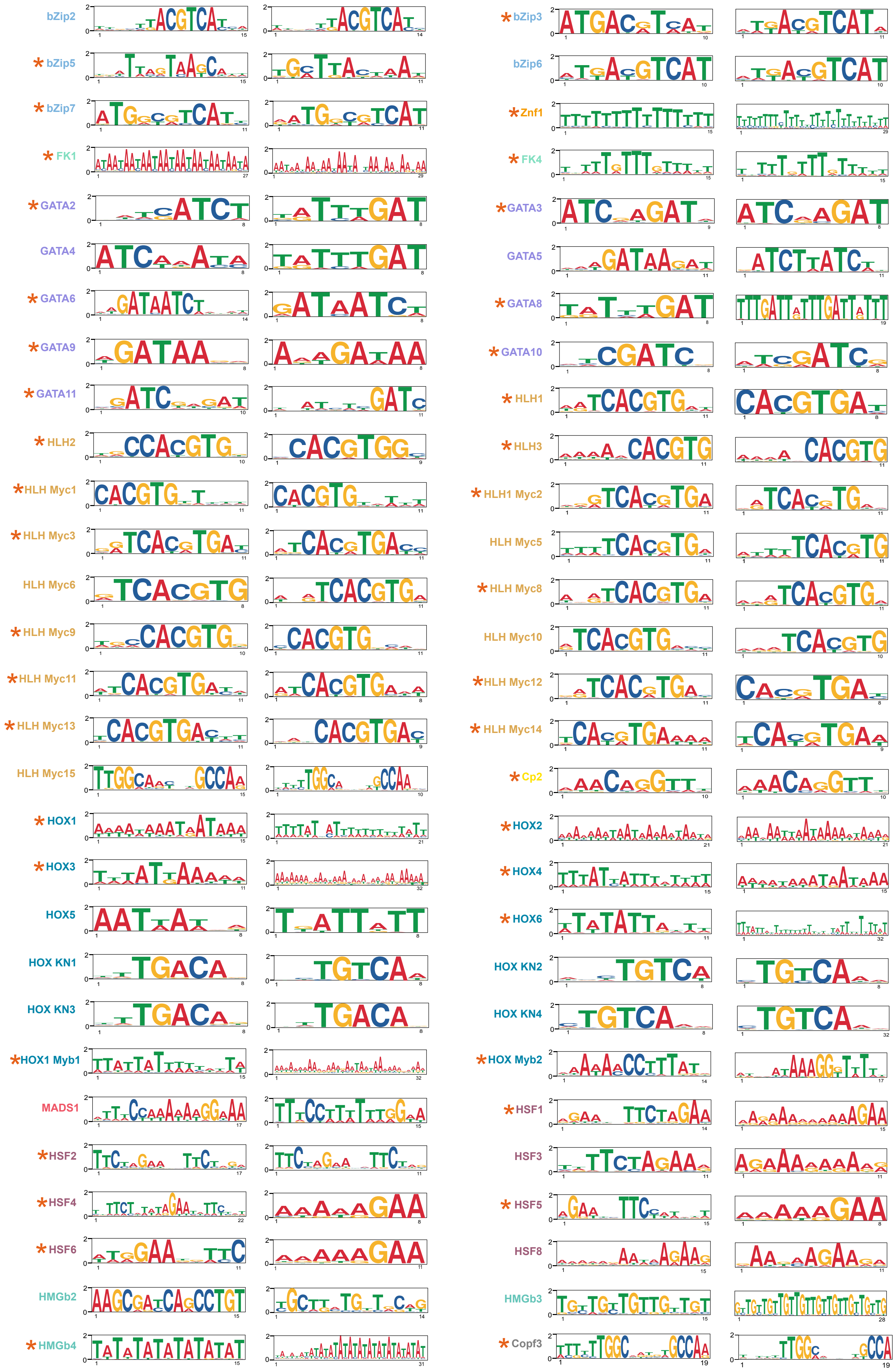
bZip2
*bZip3
*bZip5
bZip6
*bZip7
*Znf1
*FK1
*FK4
*GATA2
*GATA3
GATA4
GATA5
*GATA6
*GATA8
*GATA9
*GATA10
*GATA11
*HLH1
*HLH2
*HLH3
*HLH Myc1
*HLH1 Myc2
*HLH Myc3
HLH Myc5
HLH Myc6
*HLH Myc8
*HLH Myc9
HLH Myc10
*HLH Myc11
*HLH Myc12
*HLH Myc13
*HLH Myc14
HLH Myc15
*Cp2
*HOX1
*HOX2
*HOX3
*HOX4
HOX5
*HOX6
HOX KN1
HOX KN2
HOX KN3
HOX KN4
*HOX1 Myb1
*HOX Myb2
MADS1
*HSF1
*HSF2
HSF3
*HSF4
*HSF5
*HSF6
HSF8
HMGb2
HMGb3
*HMGb4
*Copf3

**Supplementary Figure 2.** TF motifs identified in this work. The left motifs are for the first replicate of the DAP-seq experiment, and the right motifs are for the second replicate. The orange asterisk indicates TF that passed the FRIP filter in both replicates.

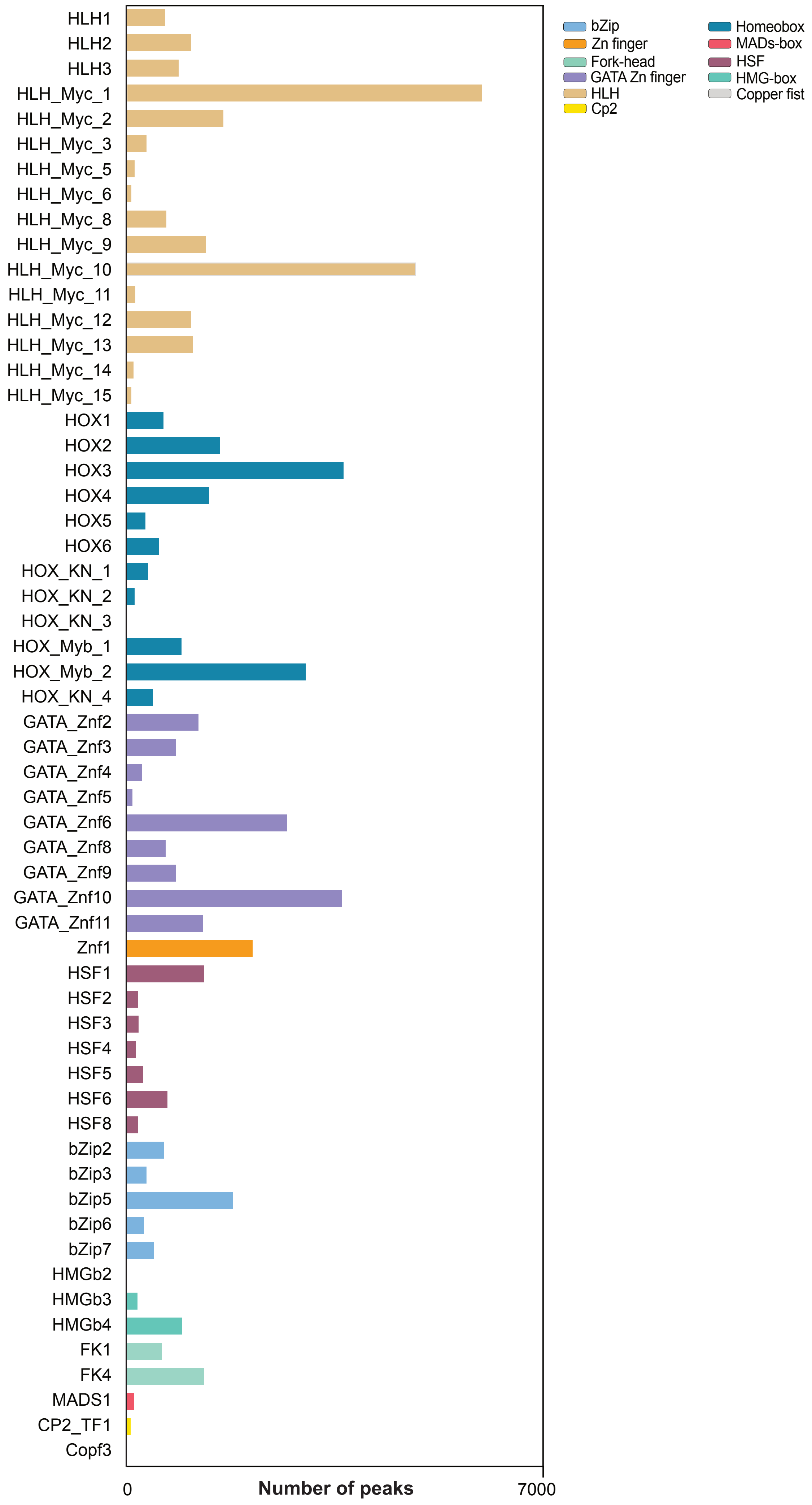
HLH1
HLH2
HLH3
HLH_Myc_1
HLH_Myc_2
HLH_Myc_3
HLH_Myc_5
HLH_Myc_6
HLH_Myc_8
HLH_Myc_9
HLH_Myc_10
HLH_Myc_11
HLH_Myc_12
HLH_Myc_13
HLH_Myc_14
HLH_Myc_15
HOX1
HOX2
HOX3
HOX4
HOX5
HOX6
HOX_KN_1
HOX_KN_2
HOX_KN_3
HOX_Myb_1
HOX_Myb_2
HOX_KN_4
GATA_Znf2
GATA_Znf3
GATA_Znf4
GATA_Znf5
GATA_Znf6
GATA_Znf8
GATA_Znf9
GATA_Znf10
GATA_Znf11
Znf1
HSF1
HSF2
HSF3
HSF4
HSF5
HSF6
HSF8
bZip2
bZip3
bZip5
bZip6
bZip7
HMGb2
HMGb3
HMGb4
FK1
FK4
MADS1
CP2_TF1
Copf3
0
Number of peaks
7000
bZip
Zn finger
Fork-head
GATA Zn finger
HLH
Cp2
Homeobox
MADs-box
HSF
HMG-box
Copper fist

**Supplementary Figure 3.** Number of peaks for each TF analyzed in this study. The different TF families are color-coded as indicated in the legend.

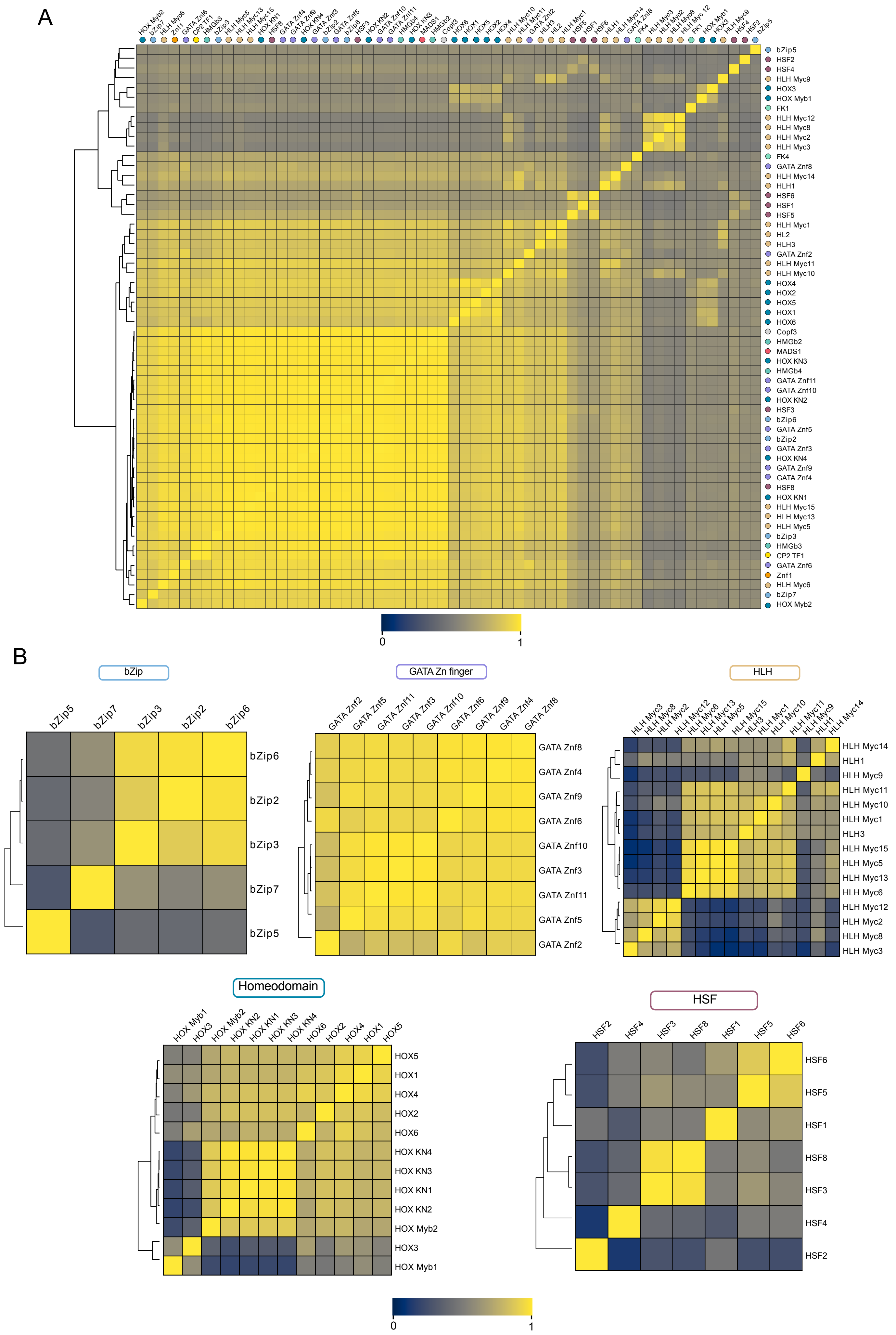
A
bZip5
HSF2
HSF4
HLH Myc9
HOX3
HOX Myb1
FK1
HLH Myc12
HLH Myc8
HLH Myc2
HLH Myc3
FK4
GATA Znf8
HLH Myc14
HLH1
HSF6
HSF1
HSF5
HLH Myc1
HL2
HLH3
GATA Znf2
HLH Myc11
HLH Myc10
HOX4
HOX2
HOX5
HOX1
HOX6
Copf3
HMGb2
MADS1
HOX KN3
HMGb4
GATA Znf11
GATA Znf10
HOX KN2
HSF3
bZip6
GATA Znf5
bZip2
GATA Znf3
HOX KN4
GATA Znf9
GATA Znf4
HSF8
HOX KN1
HLH Myc15
HLH Myc13
HLH Myc5
bZip3
HMGb3
CP2 TF1
GATA Znf6
Znf1
HLH Myc6
bZip7
HOX Myb2
0
1
B
bZip
bZip5 bZip7 bZip3 bZip2 bZip6
bZip6
bZip2
bZip3
bZip7
bZip5
GATA Zn finger
GATA Znf2 GATA Znf5 GATA Znf11 GATA Znf3 GATA Znf10 GATA Znf6 GATA Znf9 GATA Znf4 GATA Znf8
GATA Znf8
GATA Znf4
GATA Znf9
GATA Znf6
GATA Znf10
GATA Znf3
GATA Znf11
GATA Znf5
GATA Znf2
HLH
HLH Myc3 HLH Myc8 HLH Myc2 HLH Myc12 HLH Myc6 HLH Myc13 HLH Myc5 HLH Myc15 HLH3 HLH Myc1 HLH Myc10 HLH Myc11 HLH Myc9 HLH1 HLH Myc14
HLH Myc14
HLH1
HLH Myc9
HLH Myc11
HLH Myc10
HLH Myc1
HLH3
HLH Myc15
HLH Myc5
HLH Myc13
HLH Myc6
HLH Myc12
HLH Myc2
HLH Myc8
HLH Myc3
Homeodomain
HOX Myb1 HOX3 HOX Myb2 HOX KN2 HOX KN1 HOX KN3 HOX KN4 HOX6 HOX2 HOX4 HOX1 HOX5
HOX5
HOX1
HOX4
HOX2
HOX6
HOX KN4
HOX KN3
HOX KN1
HOX KN2
HOX Myb2
HOX3
HOX Myb1
HSF
HSF2 HSF4 HSF3 HSF8 HSF1 HSF5 HSF6
HSF6
HSF5
HSF1
HSF8
HSF3
HSF4
HSF2
0
1

**Supplementary Figure 4.** (A) Whole-genome coverage comparison for the 58 TF analyzed in this study. Coverage for each TF was computed in 20 kb bins. TFs were clustered according to the Pearson correlation coefficient. (B) Whole-genome coverage comparison for each specific TF family. Coverage for each TF was computed in 20 kb bins. TFs were clustered according to the Pearson correlation coefficient.

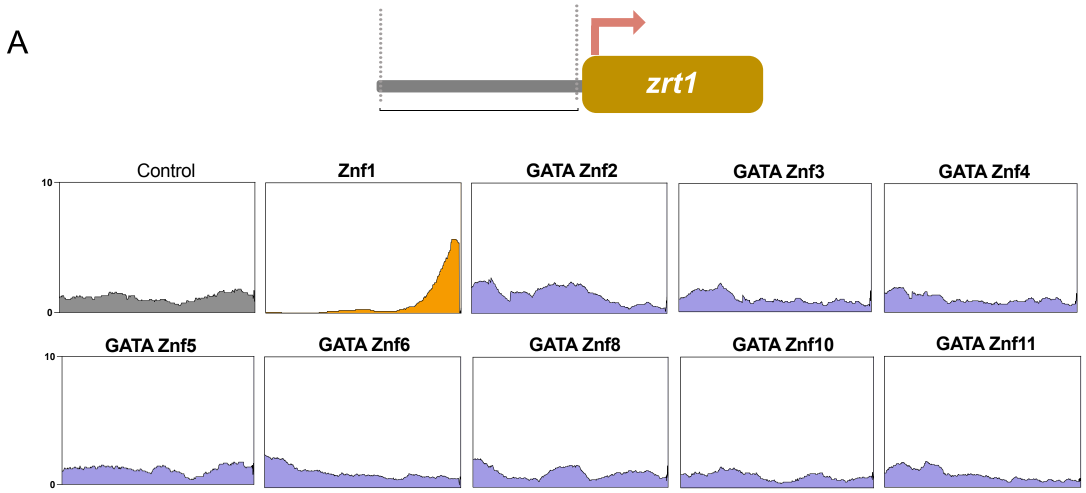


B

| | *S. cerevisiae* Protein | Rhizopus ID | log2FC (Zinc/ no Zinc) | Peak | Genomic coordinates |
|---|---|---|---|---|---|
| Experimentaly tested | ZRT1 | 1926170 | -5,61 | Yes | scaffold_17:209194-212651 |
| | ZRT2 | 1925750 | -2,14 | Yes | scaffold_17:8790-12503 |
| | ZRT3 | 1881962 | -7,65 | Yes | scaffold_2:1787719-1789186 |
| | DPP1 | | | | |
| | IZH1 | 1873310 | 0,85 | Yes | scaffold_1:546117-547577 |
| | ADH4 | 1980797 | -1,39 | Yes | scaffold_16:336523-337687 |
| | ZRC1 | 1877854 | -0,2 | Yes | scaffold_1:2692471-2694236 |
| | FET4 | | | | |
| | ZRG17 | 1872821 | -6,52 | Yes | scaffold_1:319257-320576 |
| | ZPS1 | 1888463 | -1,02 | Yes | scaffold_3:2052753-2055464 |
| Likely target | MOH1 | | | | |
| | TKL2 | 1891023 | -3,57 | Yes | scaffold_4:762883-765286 |
| | PST1 | 1911334 | -7,02 | No | scaffold_9:455146-456635 |
| | GPG1 | | | | |
| | MNT2 | | | | |
| | VEL1 | | | | |
| | COS6 | | | | |
| | MCD4 | 1898732 | -2,55 | Yes | scaffold_5:1882182-1885629 |
| | ADE17 | 1924540 | 0,93 | Yes | scaffold_15:343373-345669 |
| | URA10 | 1869696 | 1,04 | Yes | scaffold_10:307235-309190 |
| | PRC1 | 1892836 | 0,35 | Yes | scaffold_4:1556529-1557693 |
| | COS1 | 1946997 | -2,74 | Yes | scaffold_5:1375696-1392075 |
| | PHM7 | 1904003 | -0,66 | Yes | scaffold_7:296105-299332 |

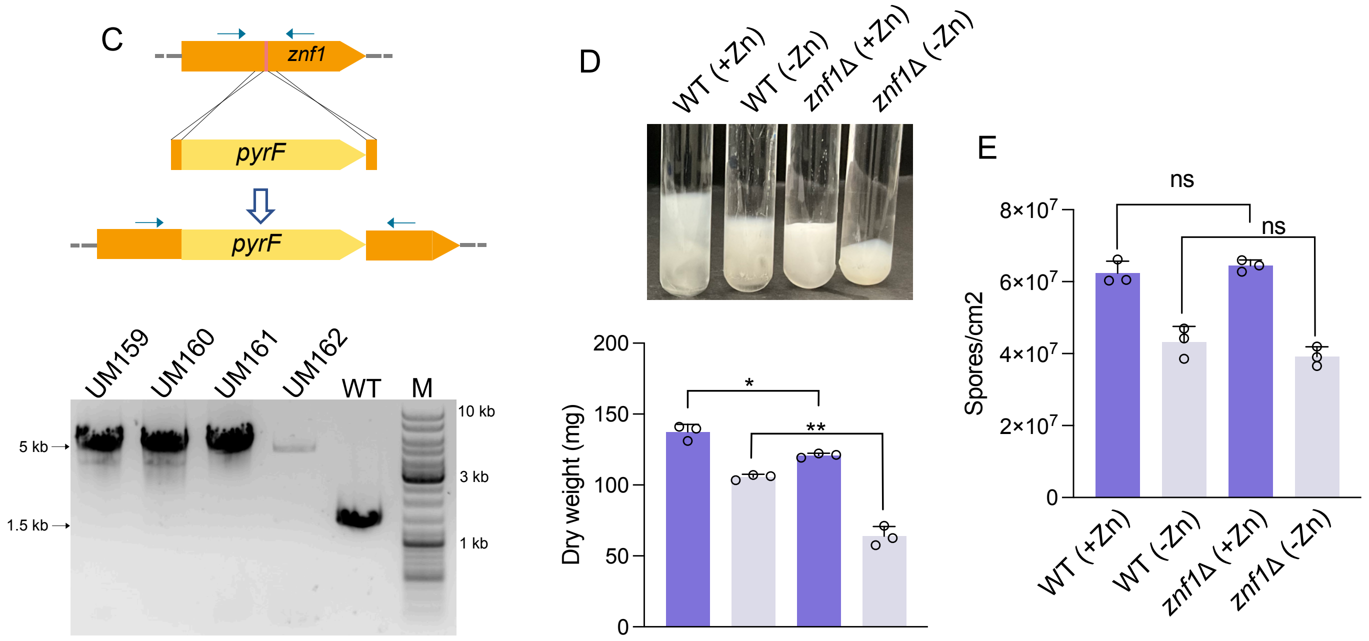

**Supplementary Figure 5.** (A) Profile coverage of all zinc finger TFs analyzed in this study over *zrt1* promoter. Coverage was plotted 1kb upstream of the *zrt1* TSS. An empty backbone plasmid was used as a control (no protein expressed). (B) Experimentally tested and likely *S. cerevisiae* Zap1 targets. The ID of the *R. microsporus* ortholog is indicated, as well as the log2FC, the genomic location, and the presence of Znf1 peak in their promoter (C) Schematic representation and results of PCR validation of the Znf1 mutant generation. Primers used to check the integration and homokaryosis are indicated as arrows in the scheme. (D) Growth of wild-type and Znf1 mutant in zinc-rich media (purple) and zinc-depleted media (gray). Image of tubes with 10 mL media is shown above the graph. A two-tailed Welch's test was performed for each comparison indicated (ns = not significant). (E) Spore production quantification wild-type and Znf1 mutant in zinc-rich media (purple) and zinc-depleted media (gray). A two-tailed Welch's test was performed for each comparison indicated (ns = not significant).

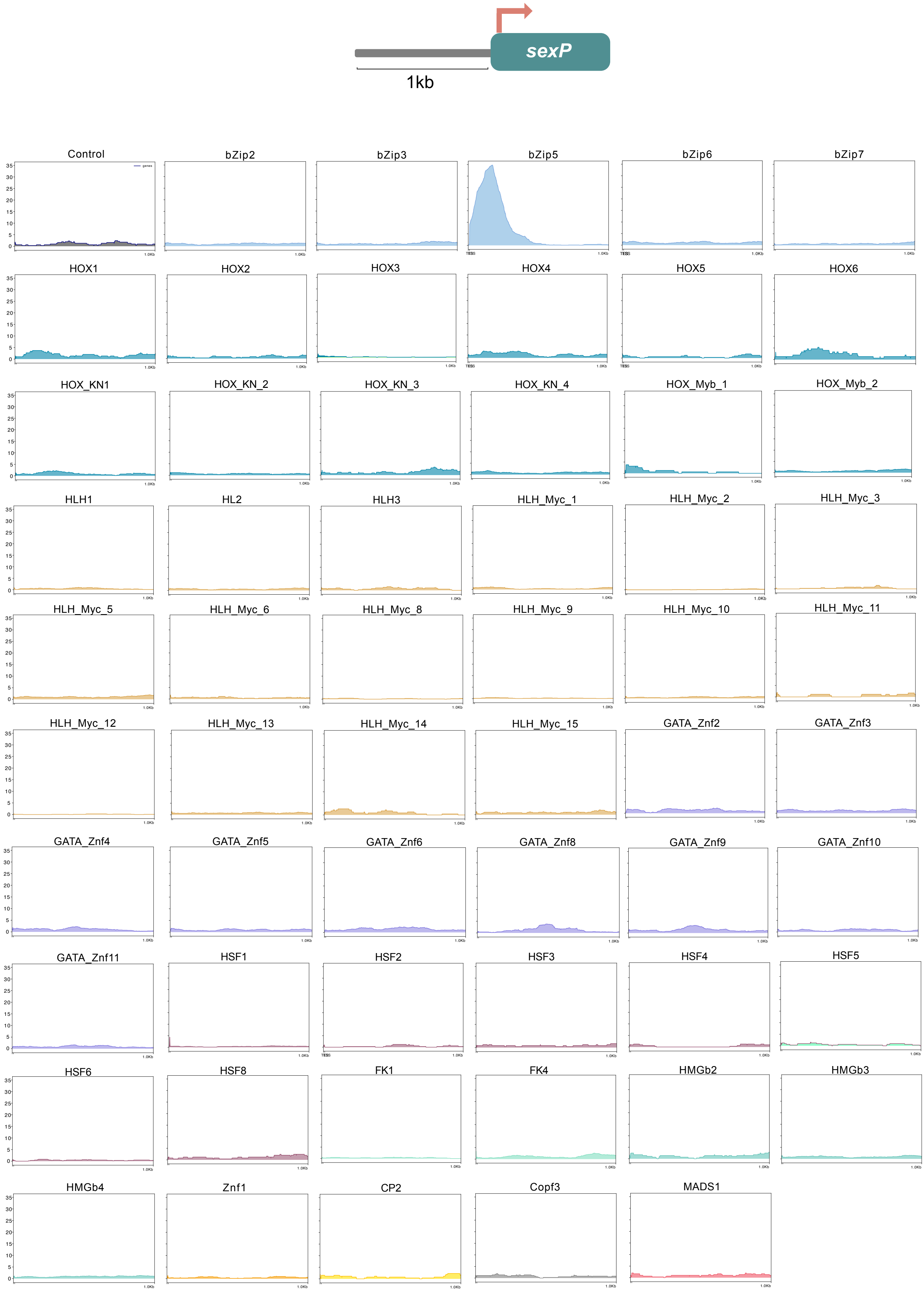
sexP
1kb
Control
genes
bZip2
bZip3
bZip5
bZip6
bZip7
HOX1
HOX2
HOX3
HOX4
HOX5
HOX6
HOX_KN1
HOX_KN_2
HOX_KN_3
HOX_KN_4
HOX_Myb_1
HOX_Myb_2
HLH1
HL2
HLH3
HLH_Myc_1
HLH_Myc_2
HLH_Myc_3
HLH_Myc_5
HLH_Myc_6
HLH_Myc_8
HLH_Myc_9
HLH_Myc_10
HLH_Myc_11
HLH_Myc_12
HLH_Myc_13
HLH_Myc_14
HLH_Myc_15
GATA_Znf2
GATA_Znf3
GATA_Znf4
GATA_Znf5
GATA_Znf6
GATA_Znf8
GATA_Znf9
GATA_Znf10
GATA_Znf11
HSF1
HSF2
HSF3
HSF4
HSF5
HSF6
HSF8
FK1
FK4
HMGb2
HMGb3
HMGb4
Znf1
CP2
Copf3
MADS1

**Supplementary Figure 6.** (A) TF profile coverage over *sexP* promoter. Coverage was plotted 1kb upstream of the *sexP* TSS. An empty backbone plasmid was used as a control (no protein expressed).

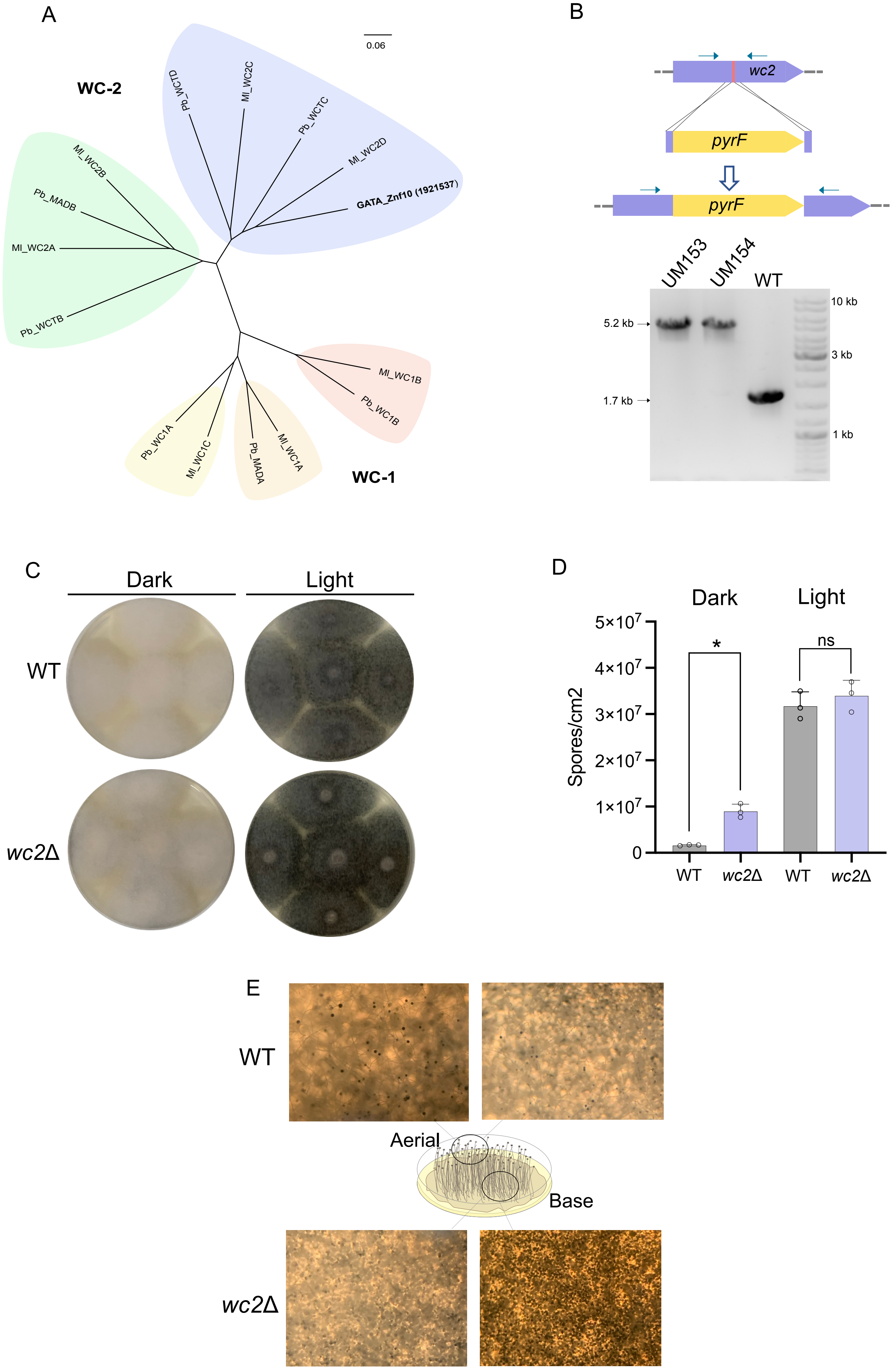
A
0.06
WC-2
Pb_WCTD
Ml_WC2C
Pb_WCTC
Ml_WC2D
GATA_Znf10 (1921537)
Ml_WC2B
Pb_MADB
Ml_WC2A
Pb_WCTB
Ml_WC1B
Pb_WC1B
Pb_WC1A
Ml_WC1C
Pb_MADA
Ml_WC1A
WC-1
B
wc2
pyrF
pyrF
UM153
UM154
WT
5.2 kb
1.7 kb
10 kb
3 kb
1 kb
C
Dark
Light
WT
wc2Δ
D
Dark
Light
Spores/cm2
5×10^7
4×10^7
3×10^7
2×10^7
1×10^7
0
*
ns
WT
wc2Δ
WT
wc2Δ
E
WT
Aerial
Base
wc2Δ

**Supplementary Figure 7.** (A) Phylogenetic tree for *M. lusitanicus* and *P. blakesleeanus* white-collar proteins with the *R. microsporus* WC-2C TF. *M. lusitanicus* and *P. blakesleeanus* known proteins were aligned with the identified *R. microsporus* WC-2C and a phylogenetic tree (neighbor-joining clustering) was generated. The distinct WC-1 and WC-2 clades are indicated in different colors. (B) Schematic representation and results of PCR validation of the WC-2C mutant generation. Primers used to check the integration and homokaryosis are indicated as arrows in the scheme. (C) Images showing *R. microsporus* wild-type (ATCC 11559) and WC-2C mutant after 48 hours of growth in the dark and under light exposure. (D) Quantification of spore production for wild-type and WC-2C mutant in dark growth and under light exposure. A two-tailed Welch's test was performed for each comparison indicated (ns = not significant). (E) Images of aerial mycelium (top) and basal mycelium (bottom) of *R. microsporus* wild-type (ATCC 11559) and WC-2C mutant. A higher sporangiophore accumulation was observed in the basal mycelium of the mutant but a lower amount in the aerial mycelium compared to the wild-type strain.

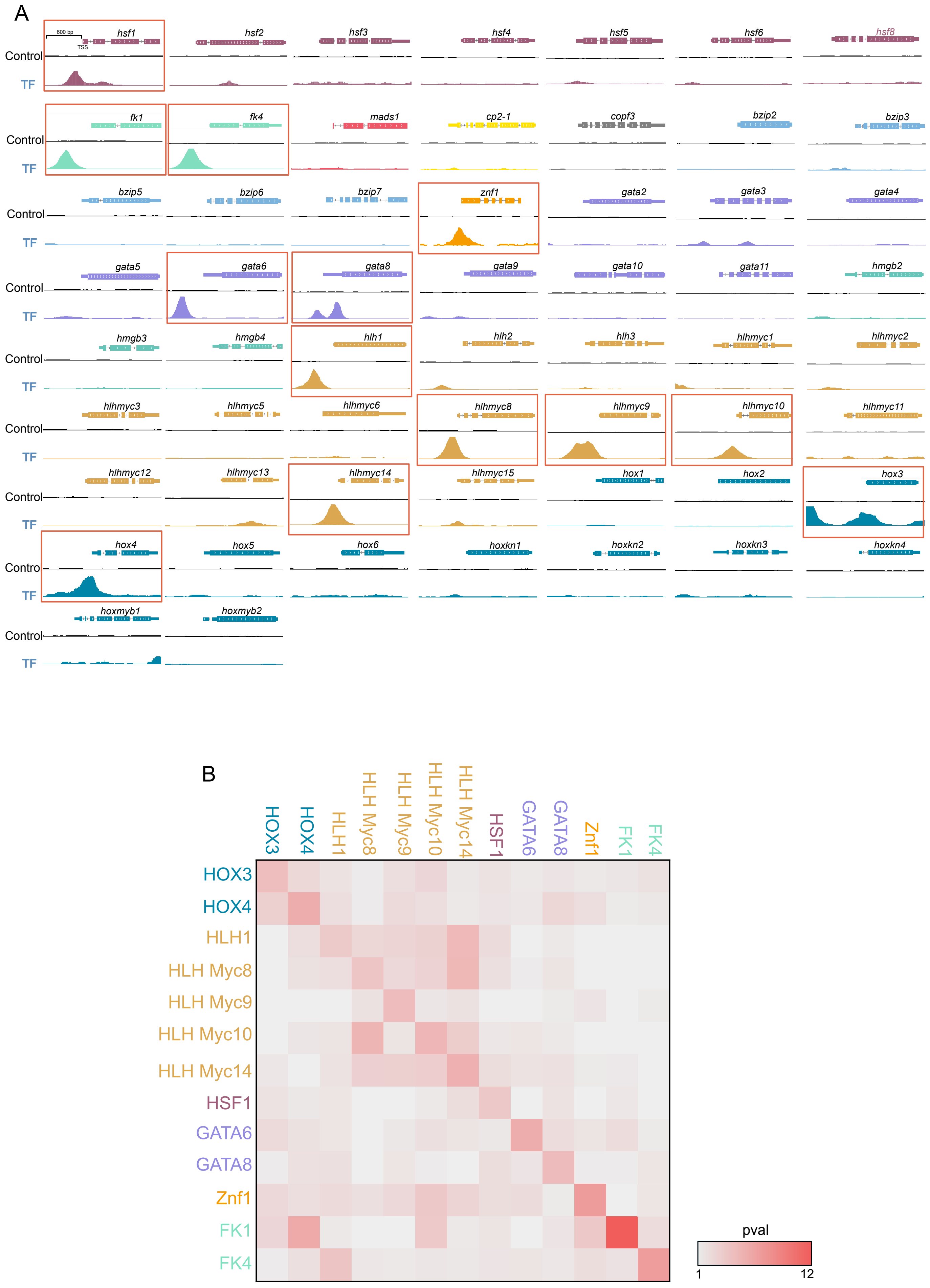
A
600 bp
TSS
Control
TF
hsf1
hsf2
hsf3
hsf4
hsf5
hsf6
hsf8
fk1
fk4
mads1
cp2-1
copf3
bzip2
bzip3
bzip5
bzip6
bzip7
znf1
gata2
gata3
gata4
gata5
gata6
gata8
gata9
gata10
gata11
hmgb2
hmgb3
hmgb4
hlh1
hlh2
hlh3
hlhmyc1
hlhmyc2
hlhmyc3
hlhmyc5
hlhmyc6
hlhmyc8
hlhmyc9
hlhmyc10
hlhmyc11
hlhmyc12
hlhmyc13
hlhmyc14
hlhmyc15
hox1
hox2
hox3
hox4
hox5
hox6
hoxkn1
hoxkn2
hoxkn3
hoxkn4
hoxmyb1
hoxmyb2
B
HOX3
HOX4
HLH1
HLH Myc8
HLH Myc9
HLH Myc10
HLH Myc14
HSF1
GATA6
GATA8
Znf1
FK1
FK4
pval
1
12

**Supplementary Figure 8.** (A) TF profile coverage over their own promoter. Coverage was plotted over the gene and 600 upstream of the TSS. Orange squares indicate TFs with a strong peak in their promoter, which are predicted to be autoregulated. (B) Motif scanning for the predicted autoregulating TFs. Motifs bound by each TF were scanned using their own promoter and the promoter of other autoregulated TFs. The significance p-val (-log10) of the best hit was used as a color scale.

**Supplementary Table 1.** Results of DAP-seq experiments. The number of peaks aligned reads and FRIP is indicated for each TF and replicate.

| JGI ID | Name | Aligned Reads (% of total) | Aligned Reads (% of total) | Peaks Exp 1 | Peaks Exp 2 | FRIP Exp 1 | FRIP Exp 2 | TF Family |
|---|---|---|---|---|---|---|---|---|
| 1762580 | HSF1 | 4589164 (94.03) | 2329209 (94.83) | 7496 | 4436 | 49.61 | 55.63 | Heat shock factor (HSF)-type |
| 1789727 | FK1 | 3638712 (73.04) | 1681306 (72.56) | 6497 | 1895 | 81.43 | 59.16 | Forkhead |
| 1796985 | Tub1 | 2596356 (91.65) | 1192264 (91.45) | 15 | 2 | 0.99 | 1.5 | Tubby |
| 1799646 | Znf1 | 4205356 (83.54) | 1362687 (87.41) | 9464 | 2650 | 76.11 | 18.38 | Zn finger C2H2/CCHC/CCCH/C5HC2 |
| 1806785 | Copf1 | 4504632 (92.77) | 2240425 (92.16) | 353 | 37 | 2.91 | 1.3 | Copper fist |
| 1832723 | bZip1 | 4517260 (92.99) | 2252529 (92.34) | 322 | 446 | 2.01 | 2.7 | bZip |
| 1867537 | MADS1 | 3637670 (73.53) | 606914 (87.02) | 3476 | 126 | 52.29 | 2.7 | MADS-box/SRF |
| 1871939 | Copf2 | 3216240 (92.92) | 2270001 (92.85) | 271 | 19 | 3.78 | 0.16 | Copper fist |
| 1872502 | CCR4_TF1 | 4490512 (92.01) | 190120 (92.28) | 30 | 0 | 0.33 | 0 | CCR4-Not complex component, Not1 |
| 1873322 | HLH1 | 4659200 (92.94) | 1679927 (89.77) | 5873 | 2622 | 49.87 | 43.17 | HLH, helix-loop-helix |
| 1873386 | HLH2 | 4603006 (93.39) | 2291532 (93.02) | 5023 | 2714 | 42.46 | 26.63 | HLH, helix-loop-helix |
| 1874682 | HOX1 | 3386712 (69.04) | 553100 (22.89) | 7928 | 1722 | 67.15 | 19.05 | HTH/Homeodomain-like |
| 1877576 | CP2_TF1 | 4222304 (84.45) | 544242 (88.45) | 4068 | 253 | 54.72 | 8.85 | CP2 |
| 1877883 | HOX2 | 2108412 (42.70) | 618797 (25.50) | 5698 | 2057 | 60.95 | 23.75 | HTH/Homeodomain-like |
| 1878704 | HOX_KN_1 | 4719012 (96.33) | 1302117 (93.12) | 4779 | 544 | 43.69 | 3.64 | HTH/Homeodomain-like (KN) |
| 1878818 | GATA_Znf1 | 851574 (92.11) | 566441 (92.16) | 35 | 3 | 0.87 | 1.28 | GATA Zn Finger |
| 1879465 | HLH3 | 4663286 (93.61) | 796714 (92.80) | 5218 | 1650 | 56.76 | 24.32 | HLH, helix-loop-helix |
| 1880317 | GATA_Znf2 | 4705120 (93.53) | 1138374 (93.20) | 9186 | 3709 | 57.15 | 31.24 | GATA Zn Finger |
| 1882186 | HLH_Myc_1 | 4571022 (93.27) | 2315726 (93.96) | 4271 | 6408 | 13.65 | 40.64 | HLH, helix-loop-helix (Myc-type) |
| 1882389 | GATA_Znf3 | 4619424 (94.11) | 2294353 (94.01) | 5254 | 1915 | 25.30 | 8.88 | GATA Zn Finger |
| 1882801 | GATA_Znf4 | 4537508 (92.16) | 2237867 (92.10) | 7189 | 664 | 36.64 | 4.35 | GATA Zn Finger |
| 1882965 | HMGb1 | 4492768 (92.42) | 2241821 (92.16) | 516 | 309 | 3.14 | 2.35 | HMG-box |
| 1884218 | HSF2 | 4554864 (93.51) | 705282 (94.22) | 4043 | 983 | 69.95 | 44.72 | Heat shock factor (HSF)-type |
| 1884678 | MADS2 | 1177878 (91.74) | 666160 (92.05) | 7 | 3 | 0.63 | 0.8 | MADS-box/SRF |
| 1884910 | Znf_PHD1 | 948074 (92.81) | 709102 (92.18) | 21 | 26 | 0.53 | 0.16 | Zn finger, PHD-type |
| 1886399 | HSF3 | 4554126 (92.73) | 476848 (91.48) | 4743 | 248 | 58.52 | 4.00 | Heat shock factor (HSF)-type |
| 1887601 | GATA_Znf5 | 4511040 (92.59) | 2261865 (92.86) | 1760 | 171 | 7.44 | 1.6 | GATA Zn Finger |
| 1887884 | HOX_KN_2 | 4455168 (93.40) | 1668154 (92.65) | 1848 | 320 | 9.74 | 2.9 | HTH/Homeodomain-like (KN) |
| 1888417 | HOX_Myb_1 | 3294632 (67.55) | 238697 (9.72) | 8249 | 1876 | 72.54 | 47.03 | HTH/Homeodomain-like (Myb) |
| 1888608 | HMGb2 | 4492432 (92.62) | 560830 (92.06) | 1057 | 45 | 5.65 | 1.39 | HMG-box |
| 1889103 | CA150 | 4503704 (92.24) | 268499 (92.70) | 388 | 17 | 1.72 | 2.03 | CA150-like |
| 1889855 | HLH_Myc_2 | 4783136 (95.70) | 2198356 (84.98) | 7442 | 2957 | 65.10 | 79.38 | HLH, helix-loop-helix (Myc-type) |
| 1890979 | HSF4 | 4396330 (90.43) | 634307 (89.14) | 6369 | 1062 | 67.63 | 35.29 | Heat shock factor (HSF)-type |
| 1891104 | HSF5 | 4518088 (92.36) | 661187 (89.92) | 5239 | 1423 | 64.85 | 25.16 | Heat shock factor (HSF)-type |
| 1891228 | HMGb3 | 4217366 (89.75) | 2109775 (89.62) | 1727 | 355 | 9.72 | 4.09 | HMG-box |
| 1891435 | HLH_Myc_3 | 4634058 (91.58) | 855288 (71.76) | 4776 | 904 | 76.04 | 53.07 | HLH, helix-loop-helix (Myc-type) |
| 1891511 | MADS3 | 287904 (91.38) | 520440 (91.86) | 0 | 3 | 0 | 1.05 | MADS-box/SRF |
| 1891808 | CBF1 | 4577342 (94.03) | 2292632 (93.94) | 218 | 4 | 1.36 | 0.35 | CBF/NF-Y/archaeal histone |
| 1892118 | HLH_Myc_4 | 397754 (91.73) | 575685 (92.06) | 58 | 18 | 1.24 | 0.15 | HLH, helix-loop-helix (Myc-type) |
| 1892227 | bZip2 | 4598570 (94.49) | 2296194 (94.18) | 4142 | 1011 | 20.61 | 4.87 | bZip |
| 1893366 | FK2 | 1051070 (91.94) | 385120 (92.08) | 9 | 6 | 0.58 | 0.24 | Forkhead/HNF3 |
| 1894049 | GATA_Znf6 | 4560906 (92.17) | 2277994 (92.31) | 7727 | 3824 | 35.68 | 21.11 | GATA Zn Finger |
| 1894449 | HLH_Myc_5 | 4556680 (92.39) | 2273168 (92.85) | 1824 | 185 | 16.44 | 1.2 | HLH, helix-loop-helix (Myc-type) |
| 1894523 | HOX3 | 4263004 (86.50) | 849154 (34.72) | 9226 | 4988 | 52.38 | 69.03 | HTH/Homeodomain-like |
| 1896330 | HLH_Myc_6 | 4441240 (90.35) | 1136250 (92.02) | 1227 | 221 | 13.99 | 3.27 | HLH, helix-loop-helix (Myc-type) |
| 1896456 | bZip3 | 4651614 (95.47) | 1740744 (93.56) | 4871 | 801 | 42.20 | 6.26 | bZip (Aft1-HRA) |
| 1896935 | bZip4 | 4556250 (93.58) | 2281740 (93.09) | 265 | 2 | 1.74 | 0.12 | bZip |
| 1897841 | HOX_KN_3 | 4187894 (96.21) | 649226 (92.51) | 2741 | 61 | 55.92 | 1.58 | HTH/Homeodomain-like (KN) |
| 1898097 | HLH_Myc_7 | 2320676 (92.14) | 356778 (91.98) | 273 | 2 | 1.99 | 0.16 | HLH, helix-loop-helix (Myc-type) |
| 1898351 | HOX4 | 3413152 (69.41) | 1027523 (42.09) | 7599 | 2642 | 54.45 | 20.09 | HTH/Homeodomain-like |
| 1898630 | HLH_Myc_8 | 4758192 (94.82) | 1905949 (75.51) | 6301 | 1242 | 68.72 | 71.42 | HLH, helix-loop-helix (Myc-type) |
| 1898716 | CoMed7 | 2476656 (91.76) | 329548 (91.96) | 7 | 20 | 0.23 | 0.16 | Coactivator Med7 |
| 1898802 | Cop15 | 4553238 (93.29) | 2281273 (93.53) | 62 | 0 | 0.18 | 0 | Coactivator p15 (PC4) |
| 1898902 | CBF2 | 1970500 (92.01) | 714157 (92.12) | 33 | 32 | 0.74 | 0.99 | CBF/NF-Y/archaeal histone |
| 1902385 | HMGb4 | 2989954 (60.31) | 1943933 (80.51) | 3009 | 1109 | 43.51 | 10.03 | HMG-box |
| 1902518 | CBF3 | 4496440 (92.09) | 284422 (91.98) | 574 | 31 | 2.81 | 0.30 | CBF/NF-Y/archaeal histone |
| 1903478 | TFIIS | 4515298 (92.74) | 490549 (92.23) | 8 | 3 | 0.73 | 1.20 | TF IIS |
| 1903516 | TFIID | 4494348 (92.15) | 435642 (92.22) | 19 | 10 | 0.26 | 1.43 | TFIID, subunit TAF12 |
| 1906134 | HLH_Myc_9 | 4631326 (93.85) | 2325476 (91.53) | 6421 | 3121 | 52.00 | 75.76 | HLH, helix-loop-helix (Myc-type) |
| 1906309 | HOX_Myb_2 | 4030268 (82.84) | 2019018 (83.00) | 5383 | 2995 | 52.60 | 30.11 | HTH/Homeodomain-like (Myb) |
| 1906563 | HLH_Myc_10 | 4543512 (92.87) | 2319200 (93.76) | 780 | 5116 | 2.57 | 36.63 | HLH, helix-loop-helix (Myc-type) |
| 1906679 | MADS4 | 670984 (91.19) | 331976 (91.82) | 13 | 6 | 0.14 | 0.92 | MADS-box/SRF |
| 1908146 | HLH_Myc_11 | 915828 (90.86) | 365485 (92.22) | 1435 | 239 | 35.78 | 6.77 | HLH, helix-loop-helix (Myc-type) |
| 1909793 | bZip5 | 4026912 (82.75) | 2101092 (87.38) | 6137 | 3644 | 76.11 | 70.33 | bZip |
| 1909873 | GATA_Znf7 | 1745278 (92.09) | 433214 (91.90) | 184 | 10 | 2.10 | 0.48 | GATA Zn Finger |
| 1909902 | HSF6 | 4571728 (93.54) | 1186297 (89.63) | 7221 | 2693 | 62.77 | 47.07 | Heat shock factor (HSF)-type |
| 1910058 | HLH_Myc_12 | 4774956 (95.49) | 2093622 (80.74) | 6051 | 1903 | 75.80 | 84.38 | HLH, helix-loop-helix (Myc-type) |
| 1914728 | HOX5 | 3484696 (71.48) | 573089 (23.82) | 7864 | 1795 | 69.58 | 18.97 | HTH/Homeodomain-like |
| 1915426 | HLH_Myc_13 | 4465516 (90.96) | 2240536 (91.52) | 2732 | 1630 | 20.10 | 9.80 | HLH, helix-loop-helix (Myc-type) |
| 1916214 | FK3 | 1103928 (91.31) | 288479 (91.94) | 21 | 1 | 0.47 | 0.18 | Forkhead |
| 1916957 | GATA_Znf8 | 4557332 (92.30) | 2068792 (91.39) | 9618 | 4193 | 47.07 | 49.76 | GATA Zn Finger |
| 1917213 | Copf3 | 4495042 (92.27) | 735766 (91.84) | 816 | 60 | 6.32 | 1.54 | Copper fist |
| 1917971 | HOX6 | 4090768 (83.84) | 296907 (34.46) | 8925 | 1218 | 58.05 | 14.49 | HTH/Homeodomain-like |
| 1918061 | GATA_Znf9 | 4509310 (92.05) | 2239214 (92.03) | 8275 | 2190 | 41.05 | 11.59 | GATA Zn Finger |
| 1919256 | HOX_KN_4 | 4623464 (94.72) | 2277772 (93.56) | 3418 | 723 | 18.90 | 4.26 | HTH/Homeodomain-like (KN) |
| 1920902 | HMGb5 | 4472412 (92.47) | 2253826 (93.17) | 58 | 59 | 1.42 | 1.34 | HMG-box |
| 1921537 | GATA_Znf10 | 4587726 (93.79) | 2263615 (93.11) | 4902 | 1580 | 21.34 | 7.17 | GATA Zn Finger |
| 1922253 | CBF4 | 694264 (91.49) | 540514 (92.12) | 43 | 11 | 0.77 | 0.07 | CBF/NF-Y/archaeal histone |
| 1923388 | bZip6 | 4599368 (94.64) | 2288313 (93.74) | 5057 | 375 | 29.40 | 2.14 | bZip |
| 1923830 | HLH_Myc_14 | 4600152 (91.89) | 454195 (93.33) | 5042 | 323 | 43.16 | 11.14 | HLH, helix-loop-helix (Myc-type) |
| 1927117 | HSF7 | 4498730 (92.79) | 2236766 (92.34) | 892 | 207 | 4.58 | 2.17 | Heat shock factor (HSF)-type |
| 1950641 | GATA_Znf11 | 4628726 (94.22) | 2263217 (93.17) | 7075 | 2404 | 31.24 | 12.64 | GATA Zn Finger |
| 1954201 | FK4 | 4037888 (81.04) | 1903070 (76.11) | 8232 | 2997 | 65.65 | 49.38 | Forkhead |
| 1958195 | HLH_Myc_15 | 4502928 (92.26) | 994869 (91.81) | 1078 | 128 | 6.40 | 0.77 | HLH, helix-loop-helix (Myc-type) |
| 1964875 | bZip7 | 4718078 (96.48) | 2270299 (93.49) | 5192 | 2169 | 64.91 | 26.28 | bZIP (HALR/MLL3) |
| 1968708 | Znf_PHD2 | 2533722 (91.49) | 427998 (92.03) | 57 | 6 | 0.56 | 0.04 | Zn finger, PHD-type |
| 1980288 | HSF8 | 4353182 (90.28) | 518446 (91.14) | 6215 | 337 | 56.49 | 2.89 | Heat shock factor (HSF)-type |
| 1982332 | HOX_Myb_3 | 4463088 (91.51) | 651390 (91.93) | 148 | 59 | 0.79 | 0.39 | HTH/Homeodomain-like (Myb) |

**Supplementary Table 2.** Total number of DE genes (upregulated and downregulated) for target genes of demethylated TFs.

| TF name | Total Targets | Total DE targets | Total Upregulated targets | Total Downregulated targets | p-val FDR Corrected Up vs Down |
|---|---|---|---|---|---|
| Znf1 | 1206 | 251 | 88 | 169 | 0,0052 (**) |
| HLH2 | 274 | 74 | 30 | 44 | 0,1599 |
| GATA10 | 1394 | 412 | 161 | 251 | 0,0454 (*) |
| GATA3 | 118 | 39 | 15 | 24 | 0,5171 |

**Supplementary Table 3.** Primers used in this study.

| Primer name | Sequence 5'→ 3' | Source |
|---|---|---|
| Znf1_locus_F | CCTACCATGCTCATCCACCT | This study |
| Znf1_locus_R | TGGCTGTCGAATCTACTTGG | This study |
| pyrF_Rm_Znf1_F | GCATGATACGAGTGAAACTTTACCAGCGCGCTCAAAGATCCTCCATAAGAATTTGACAG | This study |
| pyrF_Rm_Znf1_R | AAATGTTCATACACTTACATAAGGTTTATCTTGTCCTGTGATAAAACGAAGATGTGGCTGTC | This study |
| Rmwc2_locus_F | GTAATTCGCTGCCATGTTCA | This study |
| Rmwc2_locus_R | AGCCTCGGAGAGATCATCG | This study |
| pyrF_wc2_gRNA1_F | TCTCCAACTGGCAAAATATTGTACTGCTCAGAATCGATTCCTCCATAAGAATTTGACAG | This study |
| pyrF_wc2_gRNA1_R | CTACCAACTCATGAGGTCTATAGCCTGTTAATTCCAGTTGATAAAACGAAGATGTGGCTGTC | This study |
| A0900 | GCTCGGTAAGATGGGCTCAC | This study |
| A0901 | TGATTCTCAACTGCTCTTGC | This study |
| A1008 | GGTATCAGCAAGGCCGCAAG | This study |
| A1009 | GCTCAACTTCGTCATGTTGTACC | This study |
| A0866 | ATCGGACAACTTTCCAGACG | This study |
| A0867 | AAAGCAGGAGTTGGATGTGC | This study |
| A0854 | CGTCTGATCAATGCACAAGG | This study |
| A0855 | ACGCCCAGTTCATATTCAGG | This study |
| A0896 | TCACTCCATGCATAAGCTAGG | This study |
| A0897 | TACACCTCTCGATACACCTG | This study |
| A0856 | GCCTTAGCTTCTTTCTCTGG | This study |
| A0857 | GCCTGAAGCGCAAATGGTG | This study |
| A0870 | GATTTATCGCTCCCCTGTCG | This study |
| A0871 | TGGTCCACAGTCCTGTTCTG | This study |
| A0862 | TGGCAGAGATAGAACGTGTGG | This study |
| A0863 | TGTTGTTGTGGCTGTTGTGG | This study |
| A0167 | CGCTTCTCAAGAGCAAGACG | Lax *et al*. 2024 |
| A0168 | TGCTTGTATGGACTAATGCGATG | Lax *et al*. 2024 |
| A0894 | GTCATTCTGGTGGATTCGCC | This study |
| A0895 | ATGACCAGAAGCCTTGCCTC | This study |
| A0880 | TTTGGGCACTCGAATCAACG | This study |
| A0881 | GCCTGTGCGAGAATAGAAGC | This study |
| A0874 | ATCTTGAGAACCAGGGAGCC | This study |
| A0875 | CGGTTACGCTTGATCCTTCG | This study |
| A0914 | ATTGGTTTCGGTGGCTCCAG | This study |
| A0915 | TTCACCAGGAACAATGAAGG | This study |
| A0882 | AGTTCTGCCGCTTCTCCTAG | This study |
| A0883 | TCAACGTCGCTCAACTGATG | This study |
| A0888 | GGCTGATTGGCATGTTCGTG | This study |
| A0889 | AGTAGTAGTGTGGCCAGCAG | This study |
| A0884 | CACCATCTGAGGATTGCACAG | This study |
| A0885 | AATGTAGACTCCACCTGCGG | This study |

**Supplementary Table 4.** crRNAs used in this study.

| crRNA name | Sequence 5'→ 3' | Source |
|---|---|---|
| znf1_crRNA | TTACCAGCGCGCTCAAAGAC | This study |
| wc2_crRNA | GTACTGCTCAGAATCGATAC | This study |
